# High-accuracy structure modeling for antibody-antigen complexes

**DOI:** 10.1101/2025.11.03.686275

**Authors:** Suhui Wang, Jianan Zhuang, Xinyue Cui, Zexin Lv, Dongliang Hou, Guijun Zhang

## Abstract

Accurate modeling of antibody-antigen complexes is crucial for advancing therapeutics and diagnostics, yet predicting their binding interface remains a formidable challenge. To address this, we introduce DeepAAAssembly, a protocol that enhances static structural information from AlphaFold3 by integrating dynamic interaction patterns to guide complex assembly. Our approach leverages predicted inter-chain residue distances to construct a flexibility-aware energy function, which drives a two-stage conformational sampling process for global exploration and local exploitation. On a benchmark set of 67 representative antibody-antigen complexes, by incorporating a built-in confidence selection mechanism, DeepAAAssembly outperforms AlphaFold3, achieving not only a 12.9% higher average DockQ score and more medium- and high-quality models, but also reliably elevating the most challenging cases from incorrect to acceptable accuracy. These results demonstrate that DeepAAAssembly effectively captures conserved interaction motifs and conformational flexibility, offering a robust framework for high-accuracy antibody-antigen modeling.

## Introduction

Antibodies are fundamental components of the adaptive immune system, capable of precisely recognizing a vast array of pathogens^1^. This recognition capability, which underpins their diagnostic and therapeutic utility, stems from their structural flexibility: antibodies can reshape their complementarity-determining regions to achieve highly specific antigen binding^2^. However, this inherent flexibility also results in significant structural heterogeneity across antibody-antigen complexes, making their binding modes inherently difficult to predict. This predictive challenge is further compounded by the general lack of strong co-evolutionary signals in antibody-antigen pairs, unlike in many obligate protein complexes^3,4^. This combinatorial challenge makes the development of high-accuracy modeling methods essential for elucidating immune principles, advancing diagnostics, and guiding the rational design of therapeutics and vaccines.

The advent of AlphaFold^5^ and its successors^6–13^ has revolutionized protein structure prediction, with AlphaFold3^13^ (AF3) standing as a milestone for its high accuracy across diverse biomolecular assemblies. However, its application to antibody-antigen complexes reveals a critical gap. The reliable prediction of these complexes remains a formidable challenge, primarily due to the lack of strong co-evolutionary signals and the inherently flexible, limited-contact nature of their binding interfaces. In response, the field has pivoted towards hybrid strategies that integrate artificial intelligence (AI)-generated conformations with computational docking and experimental constraints^14–17^. Benchmarking initiatives like CASP16 confirm that such AI-augmented pipelines significantly improve model quality and confidence^18,19^. Despite these advances, the core dilemma persists: achieving a precise balance between capturing essential structural flexibility and maintaining predictive accuracy. This balance is the pivotal challenge that must be overcome to achieve robust antibody-antigen modeling.

While AI-driven methods represent the frontier of the field, traditional computational approaches to antibody-antigen complex modeling, primarily based on homology modeling and docking, continue to serve as a practical complement. Homology modeling constructs structures by reference to evolutionary templates, yet its utility is often constrained by the sequence diversity and structural flexibility of antibody complementarity-determining regions (CDRs), which limit the availability of accurate templates. Docking methods such as ZDOCK^20^ and RosettaDock^21^ probe steric and chemical complementarity through broad conformational sampling, while frameworks like HDOCK^22^ further integrate experimental data or bioinformatic priors to refine predictions. Although flexible docking algorithms^23,24^ have been developed to better accommodate binding-induced rearrangements, they still face challenges in achieving consistently high accuracy.

Notably, despite their generally lower precision, traditional docking approaches exhibit a particular strength in predicting complexes characterized by limited interfacial contact—a hallmark of many antibody-antigen interactions. This capability, combined with their ability to exhaustively sample the conformational landscape, allows them to identify plausible binding modes that might be overlooked by end-to-end AI predictors. Thus, even in the age of deep learning, such sampling-based strategies provide a valuable pathway for modeling the subtle yet challenging recognition patterns inherent to immune complexes.

In summary, AI-based prediction methods and traditional molecular docking approaches exhibit distinct complementary strengths in antibody-antigen complex structure prediction. Deep learning-based end-to-end models effectively leverage both co-evolutionary information from multiple sequence alignments (MSAs) and structural-chemical features to directly generate complex conformations that are globally structurally plausible, demonstrating exceptional prediction accuracy and efficiency when template quality is high and the interface contact area is large. In contrast, although traditional molecular docking approaches often achieve lower accuracy and rely on empirical statistical energy functions, their advantage lies in the ability to perform extensive and flexible sampling of conformational space. This characteristic enables them to perform exceptionally well in predicting complex cases characterized by small interface contact areas or substantial conformational rearrangements, allowing for the effective identification of atypical binding modes that may be overlooked by AI models. This synergy positions the hybrid strategy, which leverages AI-generated structures as initial conformations and refines them via docking-based sampling, as a highly promising approach to address the diversity and adaptability of antibody-antigen interactions, thereby overcoming key bottlenecks in complex structure prediction.

In this work, we propose DeepAAAssembly, a computational protocol that integrates deep learning-based inter-chain residue distance predictions with Monte Carlo conformational sampling to model antibody-antigen complex structures. DeepAAAssembly leverages a deep learning model, trained on physicochemical, evolutionary, and geometric features, to predict inter-chain residue distances. These predictions are combined with structural constraints from AF3 to formulate a knowledge-based energy function for guiding conformational sampling. The protocol subsequently implements a two-stage “exploration-exploitation” strategy for conformational space sampling. In the exploration stage, relative binding orientations are systematically sampled under the guidance of the energy function. In the exploitation stage, the local structure, with particular emphasis on the CDR loops, is refined to enhance geometric and energetic complementarity at the interface. The resulting conformations are evaluated using the designed energy function, which enables the selection of structurally optimal and energetically favorable models. Benchmarking demonstrates that DeepAAAssembly surpasses AF3 in DockQ and TM-score metrics by correcting systematic biases in global orientation prediction to produce more physically plausible complexes, which underscores its potential for high-fidelity antibody-antigen modeling.

## Results

### Overview of the method

DeepAAAssembly is a computational protocol designed to model antibody-antigen complex structures (**Fig. 1**). It takes the amino acid sequences of an antibody and its corresponding antigen as input, and outputs their three-dimensional complex structure. By integrating data-driven predictions of inter-chain residue distance distributions with a two-stage conformational sampling strategy, the method enables efficient exploration of flexible conformational space and accurate modeling of interface complementarity. The overall workflow consists of four main components: data preparation, feature extraction, network architecture, and a two-stage optimization procedure.

**Fig. 1.**
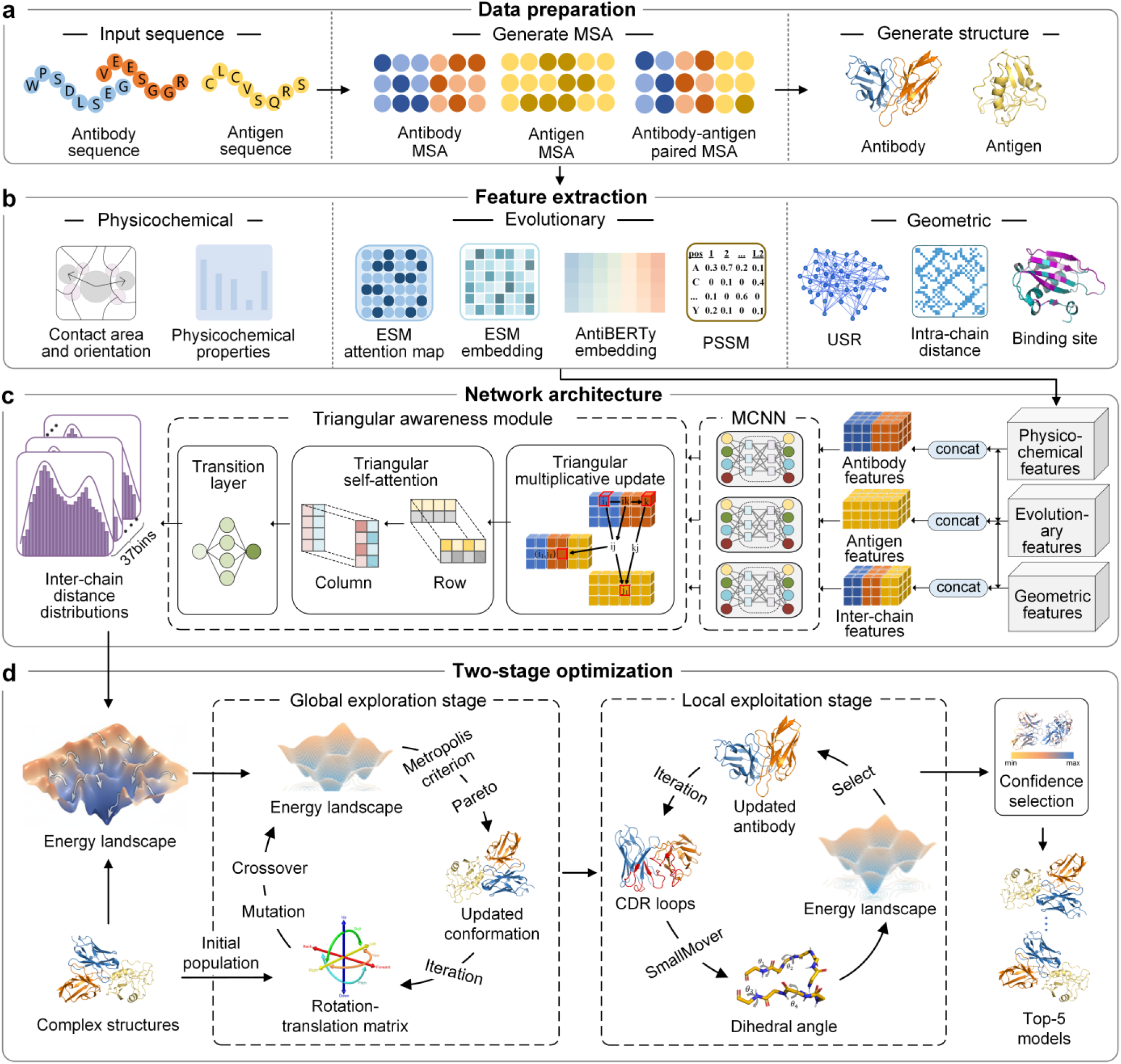
Overview of DeepAAAssembly. DeepAAAssembly takes antibody-antigen sequences as input, integrates sequence, evolutionary, and structure information to predict inter-chain residue distance distributions, and employs a two-stage conformational sampling strategy to model antibody-antigen complex structures. **a,** Data preparation. Antibody and antigen sequences are input to construct both individual MSAs for each chain and a paired MSA. Monomeric structures are generated using AF3. **b,** Feature extraction. Three complementary feature categories are extracted: physicochemical features include contact area and orientation, as well as other physicochemical properties; evolutionary features consist of ESM attention map, ESM embedding, AntiBERTy embedding, and position-specific scoring matrices (PSSM); and geometric features comprise Ultrafast Shape Recognition (USR), intra-chain distance, and binding site. **c,** Network architecture. The three categories of features are concatenated and fed into a multi-column convolutional neural network (MCNN) module, with subsequent triangular awareness module to further consider relative positional and triangular relationships between inter-chain residues, ultimately outputting inter-chain residue distance maps. **d,** Two-stage optimization. Inter-chain residue distance maps, which predicted by DeepAAAssembly and derived from AF3, jointly guide a two-stage conformation sampling process: global exploration for overall antibody-antigen orientation and local exploitation for highly flexible CDR loops. Sampled conformations are evaluated through an energy-guided confidence selection mechanism, from which the optimal conformation is identified as the final output.

During data preparation (**Fig. 1a**), MSAs and monomeric structures are generated for each chain, from which three complementary classes of features are extracted: physicochemical, evolutionary, and geometric (**Fig. 1b**). Physicochemical features include contact area and orientation^25^ as well as physicochemical properties^26^; evolutionary features comprise ESM attention map, ESM embedding^27^, AntiBERTy embedding^28^, and position-specific scoring matrices^29^ (PSSM); and geometric features encompass Ultrafast Shape Recognition^30^ (USR), intra-chain distance, and binding site^31^. Detailed description and dimensionality of the features are provided in the **Supplementary Table 1**. Among all features, several are particularly critical for characterizing distinct antibody-antigen interaction patterns. Contact area and orientation collectively quantify geometric complementarity at the binding interface, while AntiBERTy embeddings capture context-dependent residue semantics learned from large-scale antibody repertoires. This is supported by our feature relevance analysis (**Figs. 5c**, **5d**), which confirms their critical roles in characterizing interaction patterns.

In the network architecture (**Fig. 1c**), our deep network comprises two modules: a MCNN^32^ module and a triangular awareness module^33^ (Details of the network modules are provided in the **Supplementary Algorithms** section). The input features are first convolved by the MCNN module to integrate multi-scale contextual information and generate initial residue representations. These representations are then updated by the triangular awareness module, which iteratively propagates geometric information between antibody and antigen chains to enforce spatial coherence and physical plausibility. Finally, the updated representations are integrated and reprojected through the transition layer to yield refined inter-chain residue distance distributions, which are further transformed into a continuous, flexibility-aware energy landscape through probability-weighted Gaussian kernel density estimation^34^.

Guided by this energy landscape, DeepAAAssembly employs a two-stage “exploration-exploitation” optimization strategy (Details are provided in the **Methods** section) to sample conformational space (**Fig. 1d**). In the exploration stage, we introduce a multi-objective sampling strategy that integrates the classical Metropolis Monte Carlo^35^ (MMC) algorithm with Pareto dominance^36,37^, to optimize the relative orientation between the antibody and antigen. Specifically, candidate conformations are first sampled using the MMC algorithm, which accepts new conformations according to Metropolis criterion, thereby ensuring thermodynamic consistency. These candidates are subsequently evaluated via Pareto dominance analysis, simultaneously considering multiple interaction energy terms derived from distinct inter-chain distances. Only conformations that represent an optimal trade-off among these competing objectives are retained. This dual screening mechanism effectively coordinates competing interactions under multi-source distance constraints, significantly enhancing the efficiency of exploring trade-off relationships among multiple objectives within intricate binding energy landscapes. To model conformational changes induced by flexible binding, we kept the antibody framework fixed during the exploitation stage, allowing only the CDRs to undergo conformational adjustments. Specifically, we employed Rosetta’s SmallMover^38^ protocol to mitigate the high rejection rate caused by random sampling and utilized the constructed energy landscape to guide refinement, thereby generating high-quality structures with improved spatial complementarity. It is noteworthy that the energy landscape features a built-in confidence selection mechanism, where the top-ranked conformations are directly selected as final outputs based on their energy scores. Moreover, the efficacy of this mechanism has been empirically confirmed by our test results.

DeepAAAssembly establishes a rigorous protocol for modeling antibody-antigen complex structures through an integrated computational pipeline that progresses from multidimensional feature extraction to adaptive structural sampling. The protocol adaptively coordinates competing interaction terms via a multi-objective optimization strategy, thereby efficiently converging to energy-optimal conformations. The deep integration of geometric inference with this “exploration-exploitation”-based sampling strategy enables the prediction of binding modes as well as the modeling of induced-fit structural adaptations, facilitating a more comprehensive understanding of the molecular recognition mechanism.

### Modeling performance on the Docking Benchmark 5.5

To comprehensively evaluate the antibody-antigen complex modeling capability of DeepAAAssembly, we adopted the Docking Benchmark 5.5^39^ (DB5.5), which contains 67 non-redundant antibody-antigen complexes, as the test set. To mitigate potential data leakage, we filtered both training datasets to eliminate redundancy. The first training dataset, derived from the Protein Data Bank^40^ (PDB, June 2022 release) and comprising inter-chain domain pairs that we define as interacting protein domains across complex interfaces, was used for pretraining the model to capture general patterns of inter-chain interactions. After excluding entries with antibody sequences sharing more than 40%^13^ identity with those in the DB5.5 test set, this dataset comprised 10,989 inter-chain domain pairs. The second training dataset, obtained from the Structural Antibody Database^41^ (SAbDab, January 2024 release), consisted of antibody-antigen complexes and was used for fine-tuning, enabling the model to learn the distinctive physicochemical and geometric principles underlying paratope-epitope binding. Due to the high specificity of antibodies, after removing entries with antibody sequences sharing more than 99%^42^ identity, this dataset comprised 6,834 antibody-antigen complexes. Further details on dataset preparation are provided in the **Methods** section.

To systematically benchmark DeepAAAssembly against mainstream modeling approaches, we selected three representative baselines: AF3^13^, AFM^6^, and HDOCK^22^. AF3 and AFM represent the end-to-end deep learning paradigm for structure prediction, while HDOCK exemplifies the physics-based docking approach. Together, these methods represent the two dominant and complementary paradigms in the field, thus establishing a comprehensive benchmark for our evaluation. In contrast, DeepAAAssembly employs a hybrid paradigm that integrates deep learning to guide conformational sampling during the assembly process, suggesting that it may bridge the gap between the two conventional approaches. To ensure a fair comparison, each method generated five candidate structures per target. The predictions of AF3 were generated using version 3.0.0, AFM was executed using version 2.3.1, and HDOCK was run using version 1.1. All three methods were run under default parameter settings. Performance was evaluated using two complementary metrics: DockQ^43^ for assessing interface matching quality and TM-score^44^ for measuring global structural similarity.

**Figs 2a** and **2b** compare the DockQ performance of DeepAAAssembly against baseline methods for both the top1 and best models, using DockQ as our primary evaluation metric. The top1 model refers to the structure ranked first by each method’s internal confidence score, whereas the best model represents the highest-performing candidate among the five generated structures, as identified through subsequent analysis. For completeness, corresponding TM-score results for both the top1 and best models are provided in **Supplementary Fig. 1**. As shown in **Fig. 2a**, the top1 model selected by confidence from DeepAAAssembly achieves superior performance, outperforming all baseline methods in mean, median, and interquartile range DockQ values. It attains an average DockQ of 0.389, exceeding AF3 (0.366; paired Wilcoxon signed-rank test, *P* = 4.997×10^-9^), AFM (0.230; paired Wilcoxon signed-rank test, *P* = 3.528×10^-5^), and HDOCK (0.319; paired Wilcoxon signed-rank test, *P* = 1.420×10^-8^), which surpasses the state-of-the-art method, AlphaFold3, by 6.3%. We further evaluated the performance ceiling of each method by selecting the best-performing model from their five candidates (**Fig. 2b**). Under this selection scheme, DeepAAAssembly achieved an average DockQ of 0.454. This represents a significant 12.9% improvement over the state-of-the-art method, AF3 (0.402; paired Wilcoxon signed-rank test, *P* = 1.896×10^-5^). Furthermore, it substantially exceeded the performance of AlphaFold-Multimer (0.263; *P* = 2.485×10^-9^) and HDOCK (0.356; *P* = 1.026×10^-11^). These results demonstrate that DeepAAAssembly not only excels at identifying the best model but, more importantly, is capable of generating superior structural candidates. The quality of its generated ensemble suggests that its performance could be further enhanced with the development of more advanced model selection techniques^45^. A closer inspection of the score distributions reveals that DeepAAAssembly’s advantage is most evident at the median and lower quartile. In comparison, AF3 exhibits a wider interquartile range and a higher frequency of low-scoring outliers, reflecting greater performance variability. These distributional patterns indicate that DeepAAAssembly not only improves prediction reliability but also significantly reduces the likelihood of generating low-accuracy models. This consistent superiority in DockQ scores, a metric highly sensitive to interfacial contact quality and local geometric precision, further suggests that our method more accurately models the paratope-epitope interface, an advantage likely originating from its enhanced capacity to capture the intrinsic flexibility of CDRs, which undergo substantial conformational rearrangements during antigen recognition. Collectively, these results establish that DeepAAAssembly significantly enhances both the accuracy and robustness of antibody-antigen complex modeling, demonstrating a marked advance over existing deep learning and physics-based docking paradigms. Detailed results for individual targets are provided in **Supplementary Table 2.**

**Fig. 2.**
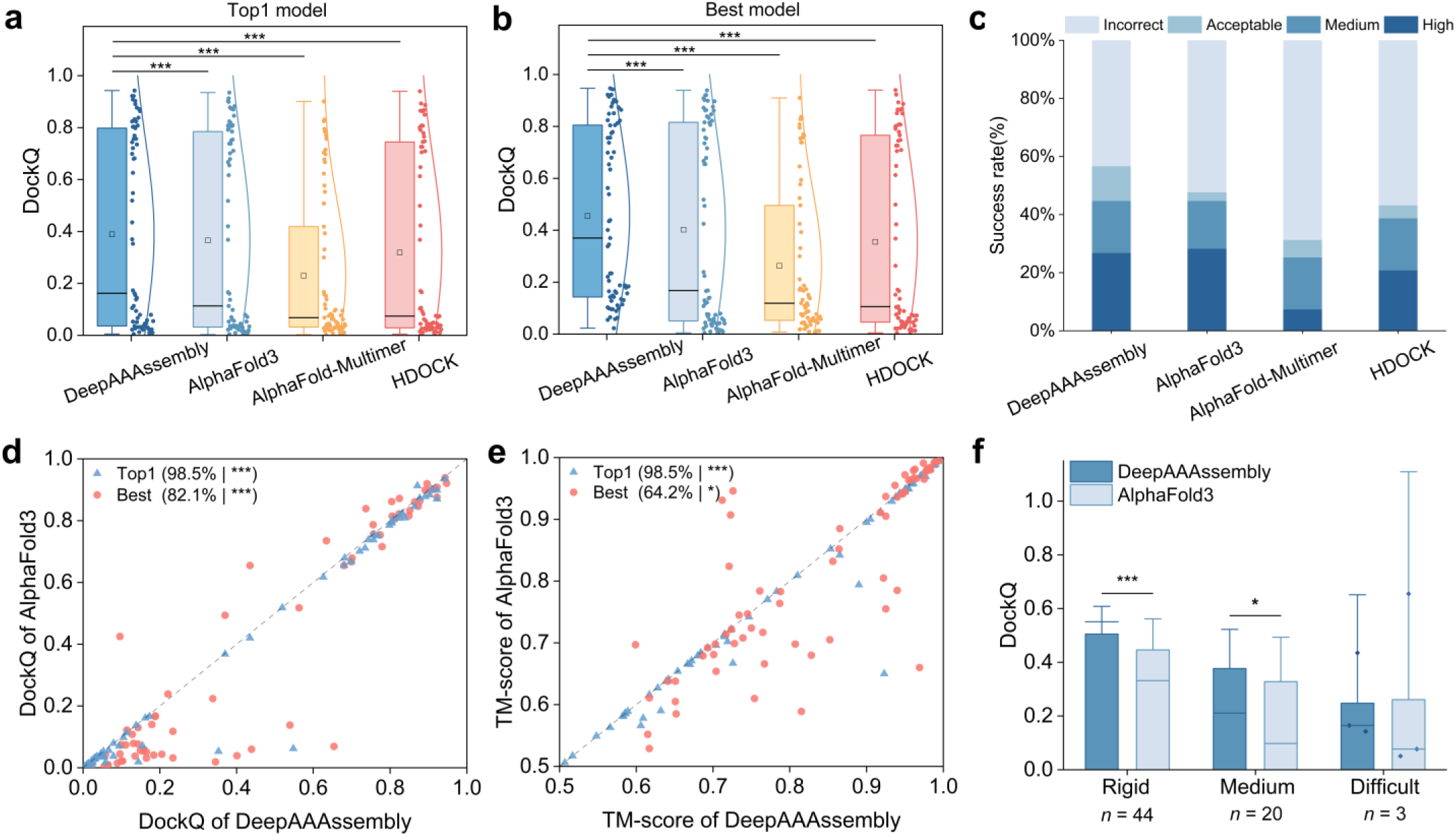
Structure modeling performance of DeepAAAssembly on the DB5.5 benchmark set. **a-b,** DockQ performance of the top1 (a) and best (b) models generated by DeepAAAssembly, AF3, AFM, and HDOCK. The box plots show the 25-75% confidence intervals (box limits), the median (centre line), the mean (square) and the range of the data distribution (whiskers). The number of asterisks denotes the statistical significance of the performance difference between DeepAAAssembly and the corresponding method. **c,** Comparison of modeling success rates among the four approaches. We followed the DockQ criteria to categorize the modeled antibody-antigen complexes into four quality levels: Incorrect (DockQ < 0.23), Acceptable (0.23 ≤ DockQ < 0.49), Medium (0.49 ≤ DockQ < 0.80), and High (DockQ ≥ 0.80). **d-e,** DockQ (d) and TM-score (e) performance comparison between DeepAAAssembly and AF3 across all test targets. The blue triangles represent the top1 models ranked by each method, while the red circles represent the best models selected based on their highest achieved DockQ or TM-score. The percentages in parentheses indicate the proportion of targets on which DeepAAAssembly achieves comparable or superior performance relative to AF3, and the number of asterisks denotes the statistical significance of the performance difference between the two methods. **f,** Performance comparison of DeepAAAssembly and AF3 in different modeling difficulty categories. *n* indicates the number of targets in each category. The number of asterisks denotes the statistical significance of the performance difference between the two methods. Due to the limited number of targets in the Difficult category, we do not report significance test statistics here. The bar height indicates the mean; error bars indicate the upper bound of the 95% confidence interval, and horizontal lines denote the medians. The statistical significance analysis above was performed by first applying the Shapiro-Wilk test to examine the normality of paired differences. Depending on the normality outcome, a two-sided paired *t*-test was used for normally distributed data, while non-normal data were evaluated using a two-sided Wilcoxon signed-rank test. Significance levels are denoted as *P* ≥ 0.05 (not significant), *P* < 0.05 (*), *P* < 0.01 (**), and *P* < 0.001 (***).

To stratify the modeled antibody-antigen complexes by quality, we leveraged the DockQ criteria^43^, categorizing them into four tiers: Incorrect (DockQ < 0.23), Acceptable (0.23 ≤ DockQ < 0.49), Medium (0.49 ≤ DockQ < 0.80), and High (DockQ ≥ 0.80). This stratification enabled a detailed analysis of performance variation across these quality groups. As shown in **Fig. 2c**, DeepAAAssembly successfully modeled 38 out of 67 targets (DockQ ≥ 0.23), achieving a success rate of 56.7%, which represents an 8.9% improvement over AF3 (47.8%). In the Acceptable quality range (0.23 ≤ DockQ < 0.49), DeepAAAssembly produced a significantly greater number of models compared to AF3, suggesting stronger robustness in accommodating the inherent conformational flexibility of antibody-antigen interfaces and in maintaining geometrically plausible binding conformations. We attribute this robustness to the fundamental design of our multi-peak distance distribution energy function (Details of the design strategy are provided in the **Methods** section). Unlike end-to-end frameworks that might converge to a single, statistically dominant conformation, modeling multiple high-probability distance states allows DeepAAAssembly to more effectively explore the conformational ensemble relevant to binding. This capability is crucial for sampling plausible interaction geometries, especially for targets where a single, rigid conformation is insufficient to describe the complex binding landscape. Beyond the Acceptable range, the comparison in higher quality tiers further highlights the competitiveness of DeepAAAssembly. In the Medium (0.49 ≤ DockQ < 0.80) and High (DockQ ≥ 0.80) group, DeepAAAssembly exhibits consistent performance with AF3 (30 targets), while substantially surpassing AlphaFold-Multimer (17 targets) and HDOCK (26 targets). These results collectively demonstrate that DeepAAAssembly maintains strong competitiveness across all quality tiers, delivering both a higher overall success rate and a more balanced distribution of accurate models. As a result, DeepAAAssembly not only advances high-accuracy antibody-antigen modeling, but also provides a more reliable baseline for acceptable conformation generation.

**Figs 2d** and **2e** present per-target scatter plots comparing the DockQ and TM-score performance between DeepAAAssembly and AF3 across individual test cases. Overall, DeepAAAssembly achieves superior performance on most targets for both the top1 and best models. Detailed comparisons of DockQ and TM-score values between DeepAAAssembly and the other methods are provided in **Supplementary Fig. 2**. In top1 model comparisons, DeepAAAssembly matched or exceeded AF3’s DockQ performance in 98.5% (paired Wilcoxon signed-rank test, *P* = 6.542×10^-9^) of targets, while maintaining comparable or higher TM-scores in 98.5% (paired Wilcoxon signed-rank test, *P* = 1.017×10^-10^) of targets. In best model comparisons, DeepAAAssembly achieved comparable or superior DockQ performance in 82.1% (paired Wilcoxon signed-rank test, *P* = 1.896×10^-5^) of targets, with better TM-scores in 64.2% (paired Wilcoxon signed-rank test, *P* = 1.559×10^-2^) of targets. The advantage of DeepAAAssembly is more pronounced in the top1 model, whereas the margin of improvement becomes smaller when the best models are compared, with the overall point distribution appearing more dispersed. The concentration of top1 data points below the diagonal indicates that DeepAAAssembly outperforms AF3 on most targets, suggesting that its modeling process consistently optimizes the input conformations. In contrast, the dispersed scatter observed in the best model plots suggests that AF3, while capable of generating highly accurate conformations, these are not always assigned the highest confidence in its internal ranking. These findings imply that DeepAAAssembly primarily enhances the reliability of its top-ranked modeling results through stable, interface-focused optimization, whereas AF3’s powerful generative framework, although capable of producing near-native conformations, does not always select them reliably. Together, these complementary characteristics reveal distinct yet synergistic strengths in conformational sampling and ranking between the two methods.

**Fig. 2f** illustrates the performance comparison between DeepAAAssembly and AF3 across different modeling difficulty categories in the DB5.5 benchmark set. According to the DB5.5 classification criteria^39^, targets were categorized as rigid, medium, and difficult. DeepAAAssembly delivers clear advantages in the rigid and medium categories, achieving average DockQ scores of 0.504 and 0.377, significantly outperforming AF3 (0.446, paired Wilcoxon signed-rank test, *P* = 1.970×10^-4^; 0.328, paired *t*-test, *P* = 3.724 ×10^-2^, respectively). For the difficult category, although the overall accuracy decreases for both methods due to the intrinsic structural complexity of these targets, DeepAAAssembly maintains performance comparable to AF3 (0.248 vs. 0.261). Notably, in two out of the three difficult cases, DeepAAAssembly achieves better results, suggesting its potential to handle challenging conformational variability. As the difficult category contains only three targets, no statistical significance analysis was performed for this group. Overall, these results indicate that DeepAAAssembly not only excels on standard cases but also preserves competitive performance on highly flexible and hard-to-model antibody-antigen complexes.

### Modeling performance on CASP antibody-antigen targets relative to AF3

To further evaluate the modeling capability of DeepAAAssembly, we constructed a benchmark set comprising 12 antibody-antigen complexes collected from the 16th and 15th Critical Assessment of Structure Prediction (CASP), including four targets from CASP16 (H1215, H1222, H1223, H1225) and eight targets from CASP15 (H1140, H1141, H1142, H1143, H1144, H1166, H1167, H1168). Notably, CASP16 was released after both of our training datasets and therefore serves as a time-separated blind set. In addition, to mitigate potential data leakage, all CASP targets from both CASP16 and CASP15 were filtered against our two training datasets using the same redundancy-removal strategy described above, ensuring that no target shared more than 40% or 99% antibody sequence identity with entries in the pretraining or fine-tuning sets, respectively.

Across the CASP16 and CASP15 tests, DeepAAAssembly and AF3 exhibit overall comparable performance in the top1 model comparison (**Fig. 3a**), and DeepAAAssembly further achieves stable improvements on the majority of medium and low-scoring samples, improving the average DockQ from 0.261 (AF3) to 0.278 (DeepAAAssembly). In contrast, when the best model is considered (**Fig. 3b**), DeepAAAssembly achieves higher DockQ scores for most targets and yields a substantial improvement in the average DockQ from 0.275 (AF3) to 0.330 (DeepAAAssembly), while successfully correcting the incorrect relative orientations of the initial conformations in multiple targets. For instance, in target H1167, the DockQ value increases from 0.130 (AF3) to 0.325 (DeepAAAssembly), exceeding the commonly accepted threshold of 0.23 and thereby moving from an incorrect to an acceptable quality range. Similarly, for H1225, the DockQ value rises from 0.159 (AF3) to 0.291 (DeepAAAssembly). These results highlight DeepAAAssembly’s ability to improve low-quality conformations, effectively transforming incorrect structures into acceptable and physically plausible models. Meanwhile, the performance gap between the top1 and best models also underscores the importance of confidence selection mechanism, indicating that DeepAAAssembly holds further potential for improvement.

**Fig. 3.**
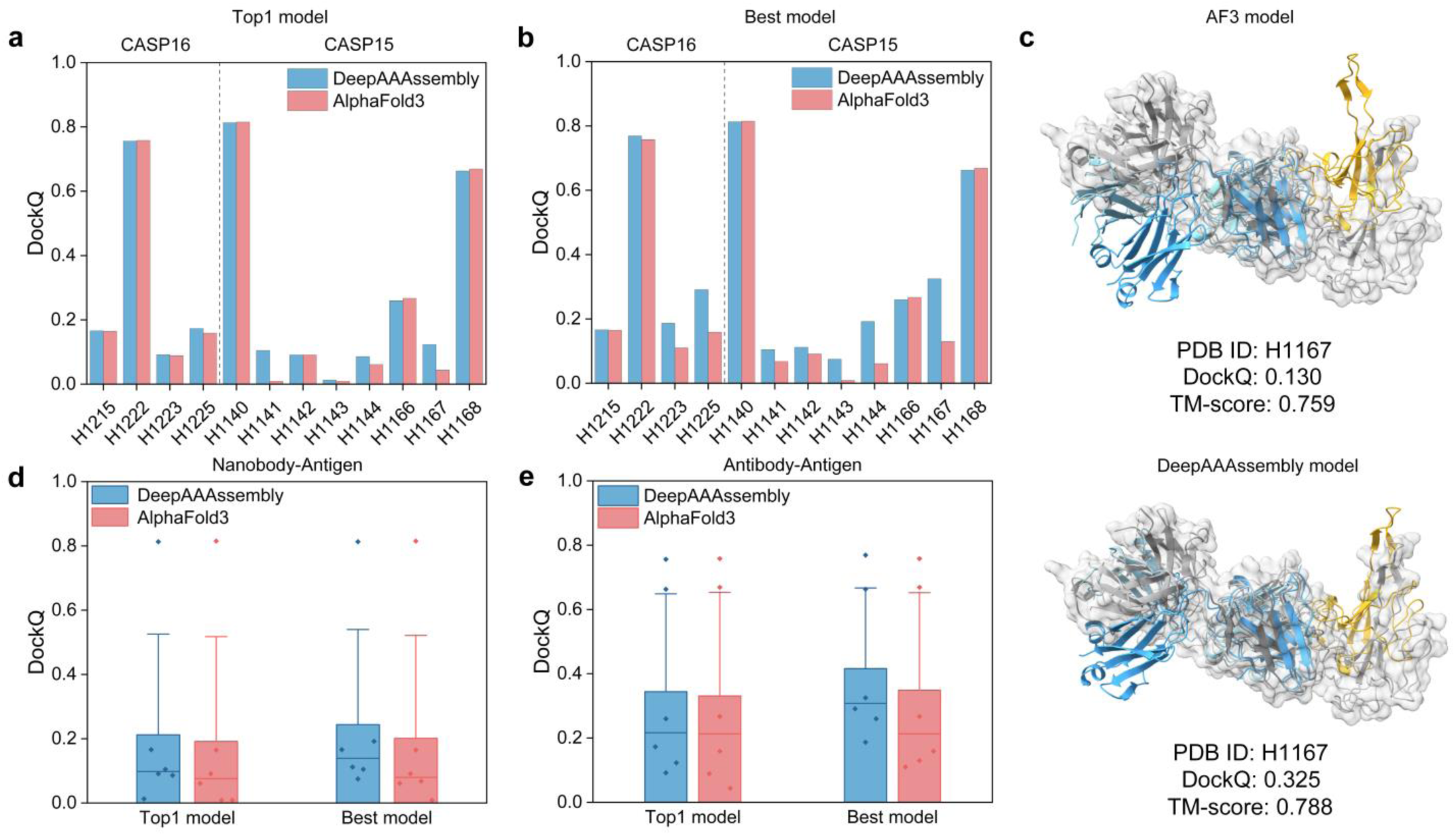
Structure modeling performance of DeepAAAssembly on the CASP benchmark set. **a-b,** Comparison of DockQ scores between DeepAAAssembly and AF3 for the top1 model and best model on each test target. **c,** Structural comparison between predicted structures and the native structure. The native structure is shown in gray, with the antibody colored in blue and the antigen in yellow. The upper figure shows the prediction results of AF3, while the lower figure shows those of DeepAAAssembly. **d-e,** Comparison of modeling performance of DeepAAAssembly and AF3 on two different antibody type complexes: Nanobody-Antigen (d) and conventional Antibody-Antigen (e). Error bars represent the upper bounds of the 95% confidence intervals for nanobody-antigen complexes and antibody-antigen complexes, respectively. Horizontal lines indicate the medians. Owing to limited dataset sizes, we do not report significance test statistics here.

To provide a more intuitive illustration, we further analyzed target H1167, whose structural models generated by AF3 and DeepAAAssembly are shown in **Fig. 3c**. The gray structure represents the experimental complex, while the antibody and antigen are colored in blue and yellow, respectively. As shown, the model generated by AF3 deviates substantially from the native structure, yielding a DockQ of 0.130 and a TM-score of 0.759. In contrast, DeepAAAssembly precisely adjusts the relative orientation between the antigen and the antibody and markedly improves the quality of the binding interface, which accounts for the substantial increase in DockQ scores.

To further investigate the impact of antibody type on modeling performance, the 12 targets were categorized into two categories: Nanobody-Antigen (H1140, H1141, H1142, H1143, H1144, H1215) (**Fig. 3d**) and conventional Antibody-Antigen (H1166, H1167, H1168, H1222, H1223, H1225) (**Fig. 3e**). Nanobody targets generally pose greater modeling challenges, as their compact single-domain architectures and highly constrained paratopes require accurate representation of localized, tightly packed interfaces. In contrast, conventional antibodies, composed of paired heavy and light chains, generally form larger and more solvent-exposed binding surfaces, which are comparatively easier to sample in structural modeling. Consistent with this structural distinction, both methods achieved lower DockQ scores on nanobody-antigen targets, reflecting the intrinsic difficulty of modeling such compact interfaces. Nevertheless, DeepAAAssembly consistently outperformed AF3 in both top1 and best-model results, and in several nanobody cases produced substantially more accurate conformations, indicating enhanced capacity to capture fine-grained interface geometry. For conventional antibody-antigen complexes, DeepAAAssembly also exhibits clear advantages, achieving higher mean and median DockQ scores, with the best-model average DockQ improved by 19.2% compared with AF3.

The analysis results suggest that DeepAAAssembly achieves consistent performance improvements across different targets. Even when the initial conformations exhibit substantial deviations in relative orientation compared with the native structures, the method can still enhance the final modeling results. This consistency reflects enhanced robustness and generalization, suggesting that the method learns transferable structural features rather than relying on case-specific patterns. These results highlight the critical role of generalization, especially in blind antibody-antigen complex modeling, where reliable performance must be achieved on unseen targets.

### Impact of predicted distance map and two-stage sampling strategy on complex modeling

To investigate the impact of our predicted inter-chain residue distance maps and the “exploration-exploitation” two-stage sampling strategy on modeling performance, we designed three ablation variants of DeepAAAssembly: w/o our distance maps (removing our predicted inter-chain residue distance maps during the conformational sampling stage); w/o exploration stage (removing the exploration stage of the conformational sampling process); and w/o exploitation stage (removing the exploitation stage of the conformational sampling process). Using the complete DeepAAAssembly pipeline as the baseline, all four configurations were evaluated on the DB5.5 benchmark set.

The ablation results of DockQ metric (**Fig. 4a**) revealed distinct performance variations across different configurations. The Baseline achieved the highest average DockQ score of 0.454 and the largest number of successful models (DockQ ≥ 0.23, n = 38). Upon removal of our predicted distance maps, the mean DockQ score decreased to 0.422 (paired Wilcoxon signed-rank test, *P* = 2.403×10^-10^) with 36 successful models. Excluding the exploration stage yielded a mean DockQ of 0.376 (paired Wilcoxon signed-rank test, *P* = 2.739×10^-11^) and 30 successful models, while omitting the exploitation stage resulted in a mean DockQ of 0.394 (paired Wilcoxon signed-rank test, *P* = 3.033×10^-12^) with 33 successful models. In particular, removing the exploration stage causes the distribution of medium- and low-quality conformations to shift significantly downward (yellow scatter points), underscoring the crucial role of this stage in elevating under-threshold (DockQ < 0.23) cases above the success cutoff.

**Fig. 4.**
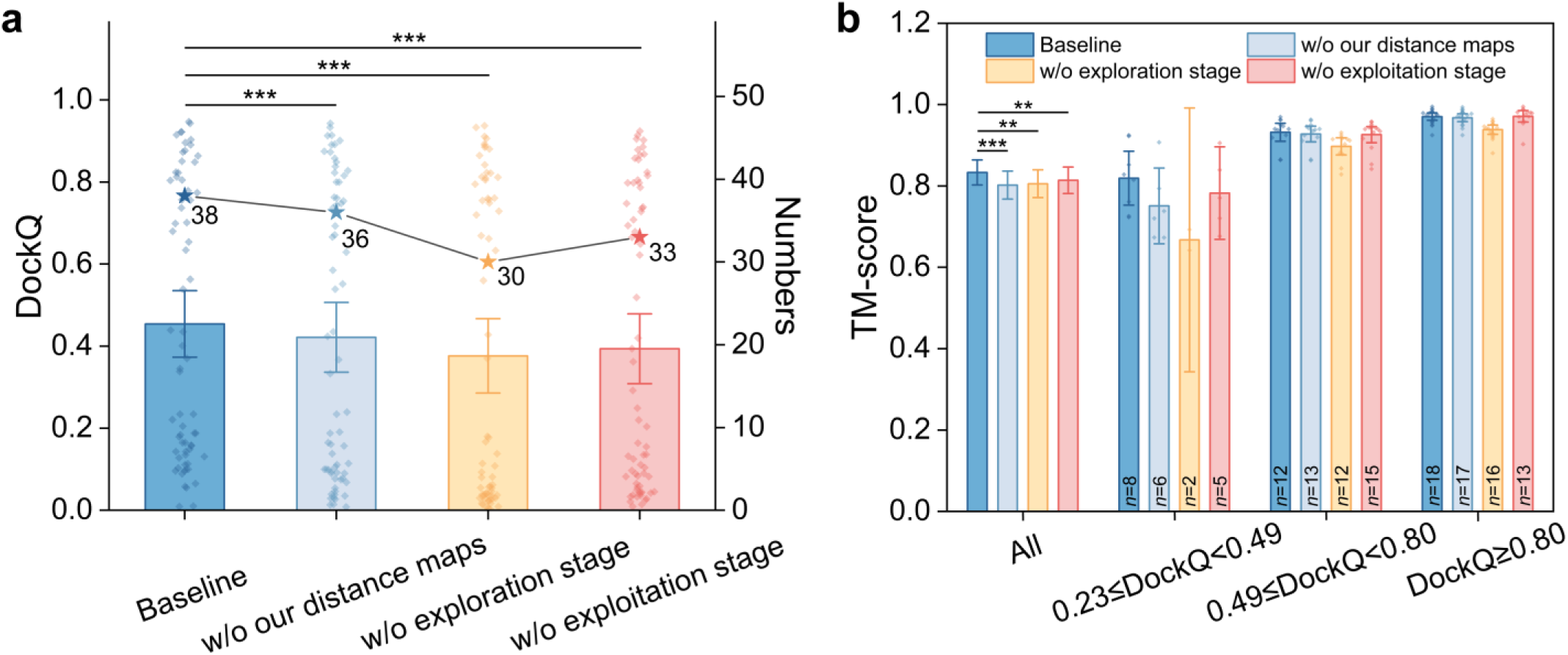
Performance comparison of different ablation variants of DeepAAAssembly on the DB5.5 benchmark set. **a,** Comparison of the mean DockQ scores on the DB5.5 benchmark set between DeepAAAssembly and its ablation variants, along with the number of targets achieving DockQ ≥ 0.23. Error bars denote the 95% confidence intervals. The number of asterisks indicates the statistical significance of the performance differences between DeepAAAssembly and each corresponding variant. **b,** Average TM-score across the full DB5.5 benchmark set and three subsets partitioned by DockQ thresholds. Error bars denote the 95% confidence intervals. The number of asterisks indicates the statistical significance of the performance differences between the Baseline and each of the three ablation variants across all DB5.5 targets. For the three ranges on the right, *n* indicates the number of targets in each DockQ range.

A more nuanced pattern (**Fig. 4b**) emerges when analyzing the TM-score results across different DockQ thresholds. When considering all samples collectively, removing our predicted distance maps led to a marked decrease in the mean TM-score from 0.833 to 0.802 (paired Wilcoxon signed-rank test, *P* = 3.357 ×10^-5^), demonstrating the essential role of accurate inter-chain distance predictions in preserving the global topology. Likewise, excluding the exploration stage and exploitation stage resulted in a reduction to 0.806 (paired Wilcoxon signed-rank test, *P* = 1.087×10^-3^) and 0.814 (paired Wilcoxon signed-rank test, *P* = 3.259 ×10^-3^), with a slightly weaker effect than that observed for the predicted distance maps. For high- and medium-quality models (DockQ ≥ 0.80 and 0.49 ≤ DockQ < 0.80), the ablation of either module exerted only a minor effect, with mean TM-scores remaining largely comparable to those of the baseline. In such cases, the conformations may already possess adequate global fidelity, thereby limiting the scope for additional improvements from these specific modules. In striking contrast, the acceptable-accuracy range (0.23 ≤ DockQ < 0.49) displayed a pronounced sensitivity to the ablations. Here, removing the exploration stage resulted in only two targets being successfully modeled, and the average TM-score dropped from 0.819 to 0.667, representing the most severe performance decline among all variants. Similarly, excluding the predicted distance maps and the exploitation stage also lowered the average TM-score to 0.751 and 0.783, respectively.

These findings indicate that the exploration stage plays a pivotal role in generating acceptable conformations, especially when guided by our predicted distance maps. We speculate that these maps capture the inherent flexibility of antibody-antigen interfaces. By encoding such probabilistic spatial information, they act as energy constraints that provide adaptive guidance during the exploration stage of conformational sampling, allowing our method to navigate complex energy landscapes and further optimize structures that are approaching their native states but have not yet reached them.

### Performance and feature analysis in inter-chain residue distance prediction

The aforementioned ablation analyses have validated the importance of our predicted distance maps for structural modeling. Building on this foundation, we select CDpred^46^ as baseline, a method widely recognized for predicting inter-residue distances in protein complexes, to perform a systematic evaluation of our deep learning predictor. The mean absolute error (MAE) is selected as the evaluation metric to measure the average deviation between the predicted and true inter-residue distances. Specifically, we evaluated the MAE for inter-chain residue distances at different cutoffs (8 Å, 12 Å, and 16 Å), where MAE is calculated only for distances falling within the respective thresholds.

Across all three distance cutoffs (**Fig. 5a**), the scatter points predominantly cluster in the upper half of the plots, significantly above the diagonal dashed line. This consistent trend indicates that for the vast majority of samples, DeepAAAssembly achieves a substantially lower MAE in predicting inter-chain distances compared to CDpred. Quantitatively, under the 8 Å cutoff, DeepAAAssembly attained an average MAE of 8.03, markedly lower than the 12.62 observed for CDPred (paired *t*-test, *P =* 1.478×10^-15^), representing a 36.4% reduction in error. At the 12 Å and 16 Å cutoffs, the average MAE of DeepAAAssembly were 5.81 and 3.91, respectively, both outperforming CDPred’s corresponding values of 9.72 (paired *t*-test, *P =* 4.694×10^-17^) and 7.03 (paired *t*-test, *P =* 2.593×10^-19^). The consistent and pronounced reduction in MAE by DeepAAAssembly’s distance maps, particularly at stricter cutoffs. It suggests that our deep learning-based approach to predict inter-chain residue distance maps is more adept at capturing the precise geometric relationships and finer details of the binding interface at close proximity. This enhanced accuracy in distance prediction for shorter-range contacts is crucial because these close-range interactions typically form the energetic hot spots of antibody-antigen interfaces, thereby playing a fundamental role in dictating binding affinity and specificity.

**Fig. 5.**
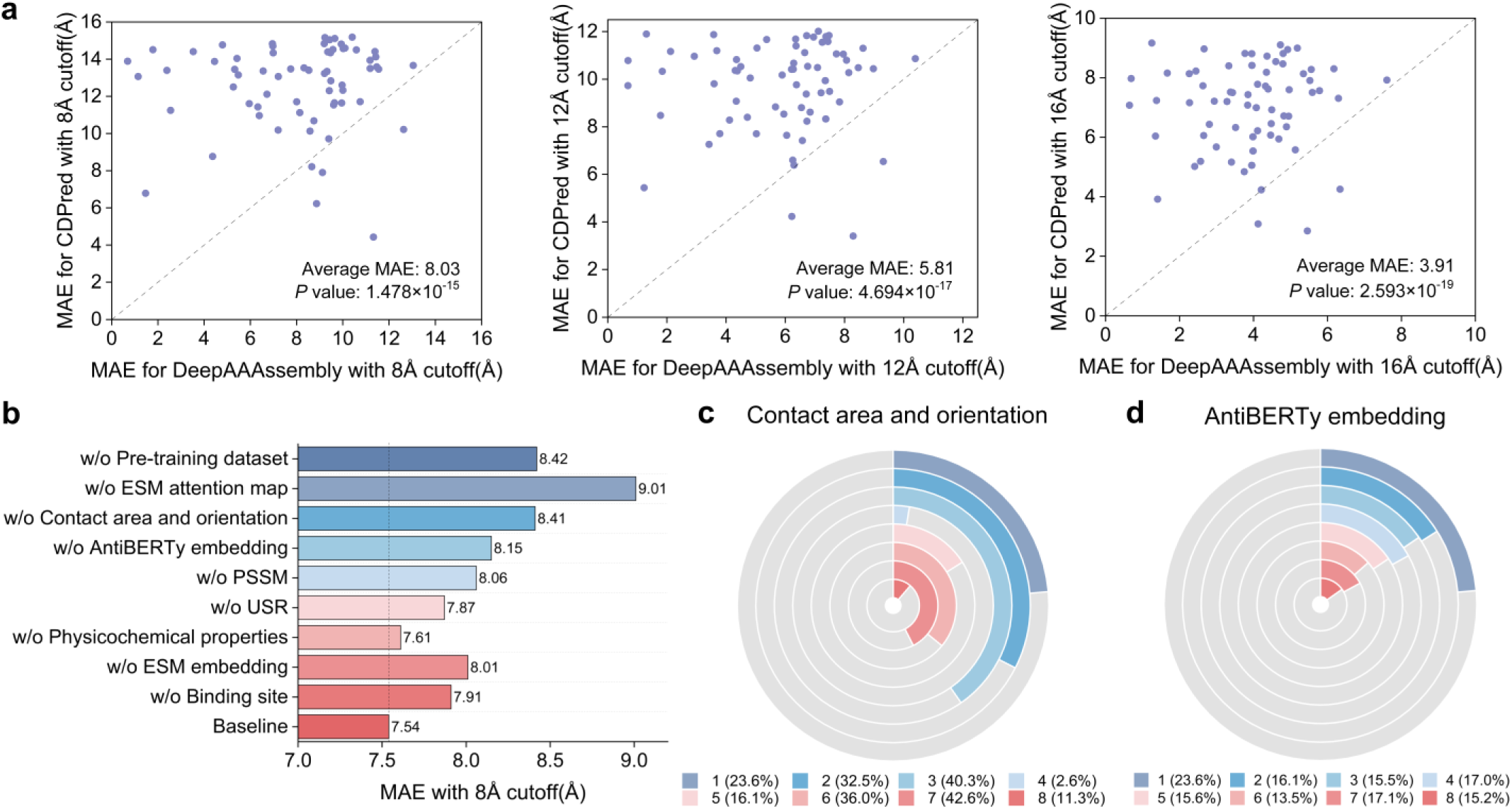
Performance comparison and feature mechanism analysis of DeepAAAssembly in the inter-chain distance prediction task. **a,** MAE comparison of inter-chain distance predictions between DeepAAAssembly and CDPred on the DB5.5 benchmark set, evaluated at distance thresholds of 8 Å, 12 Å, and 16 Å. The statistical significance (*P*-values) of the performance differences between DeepAAAssembly and CDPred at different distance thresholds is also reported. **b,** MAE performance comparison between DeepAAAssembly and its ablation variants on the DB5.5 benchmark set. **c,** Correlation between contact area and orientation features and all other features. Numbers in parentheses indicate the index of each feature: (1) AntiBERTy embedding, (2) ESM embedding, (3) ESM attention map, (4) PSSM, (5) binding site, (6) intra-chain distance, (7) USR, and (8) physicochemical properties. **d,** Correlation between AntiBERTy embedding and all other features. Numbers in parentheses indicate the index of each feature: (1) contact area and orientation, (2) ESM embedding, (3) ESM attention map, (4) PSSM, (5) binding site, (6) intra-chain distance, (7) USR, and (8) physicochemical properties.

To elucidate the factors driving the improvement in inter-chain residue distance prediction, we conducted a systematic feature ablation analysis on the DB5.5 (**Fig. 5b**). The results demonstrate that every examined feature contributes to predictive accuracy, as evidenced by the consistent increase in MAE under an 8 Å cutoff when any feature is removed. Among all tested components, the lack of a pre-training dataset led to a pronounced decline in performance, with the MAE increasing from 7.54 Å to 8.42 Å. This underscores the pivotal role of large-scale pre-training on diverse protein-protein complexes beyond antibody-antigen systems. The pre-training stage familiarizes the model with a wide spectrum of domain-domain interaction patterns from different protein complexes in the PDB, enabling it to internalize conserved geometric principles of interface complementarity, and regularities in cross-chain distances that are observed across structurally diverse systems. These conserved interaction priors can then be effectively transferred to antibody-antigen tasks, where the binding interfaces, despite their unique immunological specificity, are still constrained by the same physical principles governing protein-protein interfaces. Thus, the observed performance drop in the absence of pre-training reflects the loss of these transferable structural priors, resulting in weaker generalization and a diminished capacity to recognize plausible inter-chain geometries, particularly in regions where antigen diversity and paratope flexibility demand stronger inductive biases.

Beyond pre-training, the ESM attention maps, contact area and orientation features, and AntiBERTy embeddings exerted the strongest influences on distance prediction performance, as evidenced by the marked increase in MAE upon their removal (**Fig. 5b**). Eliminating the ESM attention maps, derived from paired antibody-antigen MSAs, caused the largest individual performance drop (MAE = 9.01 Å, 1.47 Å increase from baseline), underscoring their critical role in modeling cross-chain dependencies. Its absence substantially weakens the predictor’s ability to capture accurate cross-chain interactions and limits the triangular multiplicative update layer’s capacity to model relative positions and geometric dependencies between residues. The removal of contact area and orientation features also led to a notable decrease in accuracy (8.41 Å, 0.87 Å increase from baseline), emphasizing the importance of explicit geometric cues that encode steric complementarity and directional preferences of residue pairing, which are particularly critical for distinguishing true binding interfaces from spatially proximal but non-interacting regions. Likewise, discarding the AntiBERTy embedding caused a moderate decline (8.15 Å, 0.61 Å increase from baseline), confirming that antibody-specific contextual representations enhance the model’s understanding of paratope environments. By capturing characteristic sequence motifs and conformational flexibility patterns, these embeddings enable the predictor to identify interaction-prone loops even in the absence of strong evolutionary signals.

Furthermore, the feature relevance analysis confirmed that the selected features exhibit low redundancy. As representative examples, we selected two key features, contact area and orientation (**Fig. 5c**) and AntiBERTy embedding (**Fig. 5d**), and evaluated their correlations with the remaining features. The remaining relevant comparison results are provided in **Supplementary Fig. 3**. The results consistently demonstrate low levels of redundancy among the various features integrated into DeepAAAssembly. As shown in **Fig. 5c**, although the contact area and orientation features may exhibit some overlap with USR, the overall level of correlation across different modalities remains moderate. Similarly, **Fig. 5d** reveals that the AntiBERTy embedding also maintains relatively low correlations with features such as ESM embedding, PSSM, and particularly geometry-based features like binding site or intra-chain distance. The absence of strong, pervasive correlations suggests that each feature tends to capture unique and complementary characteristics of the antibody-antigen interface. This finding is particularly important, as it explains the robust performance observed in our ablation study: the significant performance drops upon feature removal are indeed attributable to the loss of non-redundant information channels. Thus, the integration of these diverse, low-redundancy features allows DeepAAAssembly to construct a comprehensive representation of the binding interface, encompassing sequence context, evolutionary signals, and geometric constraints, thereby enabling more accurate inter-chain residue distance prediction.

### Case study

To visually illustrate the advantages of DeepAAAssembly in modeling specific targets at both the structural and distance levels, we conducted a comparative analysis using the complex between a variant of influenza virus hemagglutinin and a neutralizing antibody binding (PDB ID: 2VIS) as a representative case. In **Fig. 6a**, the antibody and antigen are colored in blue and yellow, respectively, with the native structure shown in grey. The AF3-predicted structure exhibits a notable deviation from the native structure, with a DockQ score of 0.062 and a TM-score of 0.650. In contrast, the structure generated by DeepAAAssembly (**Fig. 6b**) achieves a substantially improved DockQ score of 0.654 and a TM-score of 0.969. Compared to AF3’s inter-chain orientation prediction errors, DeepAAAssembly more accurately captures the spatial arrangement between antibody and antigen, substantially improving both global topology and interface geometry.

**Fig. 6.**
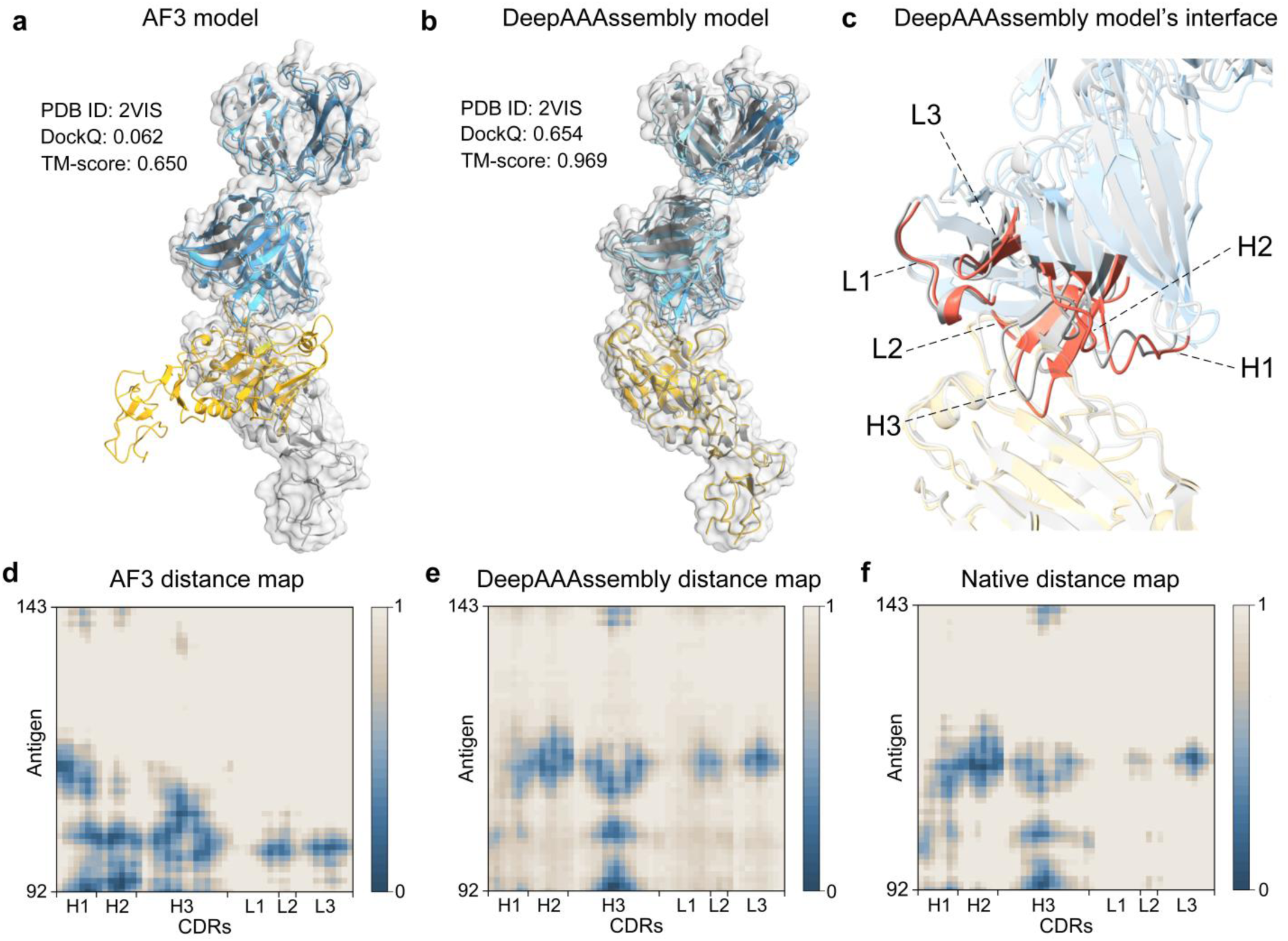
Comparative analysis of DeepAAAssembly and AF3 on the 2VIS target. **a,** Comparison between the AF3-predicted structure and the native structure. **b,** Comparison between the DeepAAAssembly-modeled structure and the native structure. **c,** Comparison of the interface region between the DeepAAAssembly-modeled structure and the native structure. **d,** Interfacial residue-residue distances extracted from the AF3-predicted structure. **e,** Interfacial residue-residue distances extracted from the distance map predicted by DeepAAAssembly. **f,** Interfacial residue-residue distances extracted from the native structure. To consistently visualize these distances, each distance matrix was linearly normalized to the range [0, 1]. The resulting normalized matrices were used to generate the heatmaps shown in Figs. d-f, where darker blue indicates shorter inter-chain distances.

Further analysis of DeepAAAssembly’s modeling performance at the interface region (**Fig. 6c**) reveals that its spatial configuration is highly consistent with the native structure. In the heavy chain, the backbone arrangements of CDRH1 and CDRH2 closely match those in the native structure. Despite the high flexibility and structural complexity of CDRH3, our method still recapitulates its backbone orientation and β-sheet characteristics to a certain extent. The conformations of CDRL1 and CDRL2 in the light chain are compact, with both the loop and α-helix structures well reproduced. The extension direction of CDRL3 toward the antigen interface and its chimeric morphology are consistent with the native structure, demonstrating that our method possesses a certain degree of spatial reconstruction capability in the key contact area.

To investigate the reasons behind the improvement in modeling accuracy, we compared the interfacial residue-pair distances extracted from the AF3-predicted structure (**Fig. 6d**), the corresponding distances predicted by DeepAAAssembly (**Fig. 6e**), and the corresponding true distances in the native structure (**Fig. 6f**). The heatmaps reveal that the distance map predicted by DeepAAAssembly closely matches the native structure in both the density and distribution pattern of contact regions, markedly outperforming AF3. This disparity highlights the critical role of accurate distance prediction in modeling interfacial residue-residue geometry. By providing fine-grained spatial constraints, the predicted distance maps effectively guide CDR conformations to better fit the antigen epitope, thereby enhancing the accuracy of the final antibody-antigen complex structure.

## Discussion

High-accuracy modeling of antibody-antigen complexes remains challenging, owing to local structural uncertainties arising from the high flexibility of CDR conformations and the strong specificity of antibody-antigen interactions. In this work, we propose DeepAAAssembly, a modeling protocol that integrates inter-chain residue distance prediction with flexible conformation sampling. This protocol employs a deep learning network to predict inter-chain residue distance maps, upon which an “exploration-exploitation” two-stage conformational sampling strategy is designed to coordinate the overall conformational search and local interface refinement, ultimately producing complex structures with global structural rationality and enhanced interface geometric quality. During sampling, multi-peak distance distributions are constructed from the predicted distance maps, integrating multiple high-probability distance peaks, which not only improves geometric adaptability but also enhances the coverage of flexible conformations. Results indicate that our predicted distance maps and two-stage sampling strategy play a synergistic role during modeling, boosting the modeling performance of AF3 in terms of both DockQ and TM-score. Notably, DeepAAAssembly not only enhances the accuracy of medium- and high-quality models but also reconstructs previously failed cases, thereby bringing them into an acceptable quality range. In particular, it models significantly more targets than AF3 within the Acceptable DockQ range (0.23-0.49) and maintains comparable or even superior performance in the Medium and High tiers, highlighting its robustness in handling flexible interfaces.

To more accurately characterize the specific interaction patterns between antibody and antigen, DeepAAAssembly incorporates a multi-source feature construction scheme and a transfer learning strategy in the task of inter-chain residue distance prediction. On the one hand, physics-aware contact area and orientation features are introduced during feature construction, systematically accounting for key physical mechanisms such as hydrophobic interactions and steric repulsion, thereby enhancing the predictor’s capacity to capture realistic interfacial contact patterns. On the other hand, the predictor is pretrained on general protein inter-chain domain dataset to extract latent conserved domain interaction patterns, providing transferable structural knowledge to antibody-antigen complexes. The integration of structural priors and transfer learning strengthens both the predictor’s representational ability for sophisticated inter-chain interactions and its generalization performance. Current mainstream methods generally do not explicitly utilize inter-domain interaction information and lack feature modeling of interfacial physical mechanisms, which limits their applicability in antibody-antigen complexes. In contrast, DeepAAAssembly achieves more accurate inter-chain distance predictions on the DB5.5 benchmark set, demonstrating its ability to capture finer patterns of inter-chain interactions and providing a more reliable geometric constraint foundation for subsequent structure modeling.

Building on DeepAAAssembly’s progress, future efforts in antibody-antigen modeling may pursue several directions. First, incorporating physics-based energy terms, such as van der Waals, electrostatic, and solvation effects, into the sampling process could improve the physical plausibility of predicted conformations. Second, embedding functional data like binding affinity into the training of distance prediction networks may help identify poses that are both structurally realistic and biologically meaningful. Finally, developing more discriminative model accuracy estimation methods specifically designed for flexible complexes would support more reliable evaluation and selection. Collectively, these directions suggest a coherent path toward structurally accurate and functionally interpretable antibody-antigen models.

## Methods

### Training set

To train a pretrained model driven by general protein inter-chain domain interactions, we collected experimentally resolved structures released before June 2022 from the PDB^40^ and applied the following filtering criteria: (1) the structure must be a protein complex containing no more than three chains, with each chain having an amino acid sequence longer than 20 residues; (2) the structure must not contain DNA or RNA molecules; (3) the experimental resolution must be better than 3.5 Å. A total of 36,955 protein complexes were retained after filtering. DomainParser^47^ was used to decompose all individual chains within the complexes into domains. Inter-chain domain pairs were then formed by combining domains from different chains within the same complex. To minimize interference from weakly interacting data during training, the interface area of each domain pair was calculated, and only pairs with an interface area exceeding 500Å^2^ were retained. Subsequently, MMseqs2^48^ was used to cluster all candidate domain pairs against the antibody portions of targets in our test set at a 40%^13^ sequence identity threshold, and all entries belonging to the same cluster as any test target were removed. After redundancy removal, 16,336 inter-chain domain pairs remained. To meet the needs of the model task, only 10,989 heteromeric domain pairs were used as the inter-chain domain dataset. These were then randomly split into a training set (9,890 pairs) and a validation set (1,099 pairs) in a 9:1 ratio.

To train a distance predictor driven by antibody-antigen inter-chain interactions, we collected antibody-antigen complexes released before January 23, 2024 from the SAbDab^41^. The IMGT^49^ numbering scheme was adopted, and the following criteria were used for filtering: (1) experimental resolution better than 4.5 Å; (2) antigen chains are present and of protein type; (3) the antibody must contain at least a heavy chain. Based on the annotations provided by SAbDab, the antibody-antigen complex was separated into complex structures in which the antibody chain and the antigen chain were paired one by one, and 7014 antibody-antigen complex structures were obtained. MMseqs2^48^ was subsequently employed to cluster the antibody sequences in the candidate data with those of the targets in our test sets at a 99%^42^ sequence identity threshold, and removed all entries belonging to the same clusters as the test targets. A relatively high redundancy threshold (99%) was adopted because antibody specificity is largely governed by highly localized hypervariable regions (CDRs); applying a more stringent threshold would eliminate many entries that differ in functionally relevant CDR segments despite sharing similar framework sequences. This process resulted in 6,834 non-redundant antibody-antigen complex structures. The dataset was randomly divided into training and validation sets in a 9:1 ratio. The training set comprised 6,131 samples, and the validation set contained 703 samples.

### Test set

To evaluate the performance of antibody-antigen complex structure modeling methods, we selected the DB5.5 benchmark set as the primary test set, which comprises 67 non-redundant antibody-antigen complex structures. Additionally, we curated 12 available antibody-antigen complex targets from the CASP16 and CASP15 datasets to form a CASP benchmark set. The experimentally resolved structures of all targets can be obtained in the PDB, providing a reliable benchmark for performance evaluation. These targets are listed under the PROTEIN MULTIMERS category for CASP16 (https://predictioncenter.org/casp16/) and the MULTIMERS category for CASP15 (https://predictioncenter.org/casp15/) on the official CASP websites.

### Features extraction

Prior to feature construction, we generated MSAs, paired MSAs, and corresponding structures for both antibody and antigen input sequences. Although antibody-antigen complexes generally lack pronounced co-evolutionary relationships in nature, we still observed that weak co-evolutionary relationships do exist, often manifested in the relatively shallow depth of the paired MSAs. Therefore, we consider that paired MSAs may contain statistical features reflecting structural complementarity or species co-occurrence patterns^50^. Such information can provide implicit structural constraints for deep learning model, facilitating the identification of physicochemical complementarity (e.g., shape, charge, and hydrophobic matching at the interface) or functional relevance (e.g., species-specific pairing or evolutionary co-occurrence) between antibody and antigen. For samples containing both antibody heavy and light chains, we maintained independent molecular representations in the predictor to avoid feature contamination caused by directly concatenating the two chains. The same independent processing strategy was adopted during the construction of antibody MSAs and antibody-related features. Monomeric MSAs were generated by searching the UniRef30 (2021-03) database^51^ using HHblits^52^. The specific parameters are listed in **Supplementary Table 3**. Paired MSAs were constructed based on species annotation information by selecting and concatenating sequence pairs from antibody and antigen MSAs sharing identical species annotations. For antibody samples containing both heavy and light chains, the antibody MSAs were constructed by concatenating heavy and light chain MSAs according to species annotations. During model inference, antibody and antigen structures were predicted using the AlphaFold3 with default parameter settings.

The features used in the distance predictor mainly include three categories: physicochemical features, evolutionary features, and geometric features. Physicochemical features include the physics-aware contact area and orientation features^25^, as well as amino acid physicochemical properties^26^ such as isoelectric point, polarity, and secondary structure. Evolutionary features include attention maps and vector embeddings generated by the ESM-MSA-1b language model^27^, sequence embeddings from the antibody-specific pre-trained model AntiBERTy^28^, and PSSM^29^. Geometric features comprise USR^30^, intra-chain residue distances, and binding site^31^. Among all features, the contact area and orientation features are particularly critical. By quantifying the inter-residue contact area, solvent-accessible surface area, and spatial orientation of contacting residues, they characterize the strength of physical interactions between antibody-antigen surface regions and the surrounding solvent, as well as the spatial relationships between surface and buried residues of the antibody and antigen. This characterization not only facilitates the identification of potential interface regions and reveals energetic coupling relationships, but also compensates for the limitations of sequence-based features in capturing fine-grained interaction patterns, thereby providing physically meaningful guidance for modeling complex interface conformations. A detailed description of the contact area and orientation features is provided in **Supplementary Note 1**. Comprehensive information on all input features can be found in **Supplementary Table 1**.

### Network architecture

The network architecture consists of a feature preprocessing module, a multi-column convolutional neural network^32^ module, and a triangular awareness module. The raw input features comprise physicochemical, evolutionary, and geometric features. In the feature preprocessing module, these features are reorganized into antibody features, antigen features, and inter-chain features. All one-dimensional features are first cross-concatenated in the feature preprocessing module to expand them into two-dimensional feature tensors suitable for convolutional operations. Specifically, the one-dimensional features are duplicated along the sequence-length dimension and concatenated along the channel dimension to generate antibody and antigen self-cross feature tensors, as well as antibody-antigen cross feature tensors. Detailed algorithms are described in **Supplementary Algorithm 1**. All resulting two-dimensional features are processed through a 1×1 convolutional layer to unify the number of channels to 64, followed by instance normalization and subsequent activation using a leaky rectified linear unit (Leaky ReLU). Finally, the processed two-dimensional features are concatenated by categories (antibody, antigen, and inter-chain features), and subjected to dimensionality reduction, normalization, and activation once more to produce a unified input tensor for the subsequent modules.

The MCNN module comprises three parallel convolutional branches, each employing convolutional kernels of different sizes to enable multi-scale feature extraction. The first branch utilizes 9×9 and 7×7 kernels, the second employs 7×7 and 5×5 kernels, and the third adopts 5×5 and 3×3 kernels. By integrating convolutional kernels of varying scales, the network is capable of capturing spatial relationships and structural dependencies among residues at both local and global levels. Large kernels enhance the ability to model long-range structural information, while smaller kernels contribute to the characterization of local topological relationships. To strengthen the network’s ability to represent features under diverse inter-chain interactions, instance normalization is used in place of batch normalization. Additionally, residual connections are incorporated into each convolutional branch to alleviate the vanishing gradient problem and improve cross-layer feature integration capability. Detailed implementation is provided in **Supplementary Algorithm 2**.

To further enhance the predictor’s capacity for geometric relation modeling, we incorporate the triangular awareness module originally proposed in AlphaFold2^5^, and inspired by the DeepInter^33^ method, extend it from intra-chain to inter-chain applications to improve the accuracy of inter-chain residue distance prediction. This module consists of four sub-layers: a triangular multiplicative update layer, a row-wise triangular self-attention layer, a column-wise triangular self-attention layer, and a transition layer. The triangular multiplicative update layer constrains the pairwise representation to satisfy the triangle inequality based on the geometric relationships among residue triplets. In the row-wise triangular self-attention, for each antibody residue (a fixed row), the interactions with all antigen residues are updated jointly via attention, allowing information flow among all pair representations that share the same antibody residue. Symmetrically, in the column-wise triangular self-attention, for each antigen residue (a fixed column), the interactions with all antibody residues are updated, enabling information flow among all pair representations that share the same antigen residue and dynamically emphasizing critical inter-residue interactions Detailed algorithms are provided in **Supplementary Algorithms 3 and 4**. This design enables the predictor to extract interaction signals bidirectionally and strengthens its capacity for context-dependent modeling. Finally, the transition layer is composed of two fully connected layers to adjust feature dimension and enhance nonlinear representational power.

### Transfer learning

To mitigate overfitting risks arising from the limited training samples of antibody-antigen complexes, we adopt a data-driven transfer learning strategy that transfers knowledge learned from inter-chain domain dataset to the antibody-antigen distance prediction task. Specifically, the predictor is first pre-trained on an inter-chain domain dataset constructed from heterologous protein complexes, aiming to learn generalizable representations of inter-residue contacts. During this phase, the input to the model excludes any antibody-specific prior information and relies solely on general one-dimensional and two-dimensional protein features, with the goal of capturing transferable spatial geometric patterns.

Subsequently, the pre-trained model is fine-tuned on the antibody-antigen complex dataset. To fully leverage antibody-specific sequence-structure information, antibody sequence embedding features generated by the AntiBERTy pre-trained model are incorporated during the fine-tuning phase, with corresponding adjustments made to the channel dimension to ensure consistent feature fusion. This transfer process helps to enhance the predictor’s capacity to capture antibody-specific residue distributions and the flexible conformations of variable regions (e.g., CDR loops), thereby boosting the performance of inter-chain residue distance predictions.

To strengthen the predictor’s discriminative ability for antibody-antigen contact regions, a class difficulty-weighted focal loss function^53^ is introduced, formulated as follows:

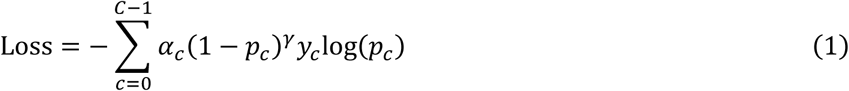

where *C* denotes the number of bins used to discretize the predicted distances, set to 37; *α*_*c*_ represents the weight of the *c*-th bin; *p*_*c*_ denotes the predicted probability on the *c*-th bin; *γ* is the focusing parameter set to 2; *y*_*c*_ is the one-hot encoding value of the true label on the corresponding bin. This loss function is designed to mitigate the imbalance in the distribution of inter-chain residue distance labels. The specific settings for the bin weights are provided in **Supplementary Table 4**.

Model training was conducted using the Adam optimizer with an initial learning rate of 0.0005, which was gradually reduced following an exponential decay strategy with a decay rate of 0.99. The training was performed for 25 epochs with a batch size of 1. All training procedures were carried out on an NVIDIA A100 GPU, with a total runtime of approximately eight days.

### Global conformation exploration through multi-population multi-objective optimization

After obtaining our discrete inter-chain distance distributions, we further incorporated the distance maps extracted from the top five AF3-predicted structures to construct geometric restraints, which were then used to guide conformational sampling. Specifically, for the discrete inter-chain distance distributions, we first adopt a Gaussian kernel density estimation^54^ (KDE) to fit the 37 discrete distance bins into a continuous, multi-peak probability distribution, with the x-axis representing the continuous distance values and the y-axis denoting the probability density of residue pairs at the corresponding distance. To more finely characterize the geometric preferences within the structure, we further extracted peaks with distances less than 12 Å as spatial priors information, aiming to retain more potential local optima solutions and enhance the capacity to represent conformational diversity. During the sampling process, we calculated the Euclidean distance 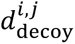 between the C_β_ atoms of antibody-antigen residue pairs in each candidate conformation and evaluated the deviation from each peak distance 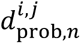 in the probability distribution. The weights were determined by the normalized probability density corresponding to each peak. The energy function is defined as follows:

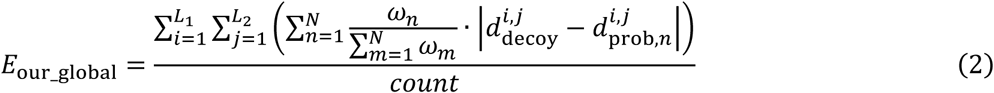

where *L*_1_ and *L*_2_ denote the sequence lengths of the antibody chain and antigen chain respectively. *ω*_*n*_ represents the probability density value corresponding to the distance of the *n*-th peak, *N* indicates the number of peaks within 12 Å for a given residue pair, and *count* refers to the number of residue pairs with distances greater than zero.

Correspondingly, we directly extract the C_β_ atom distances 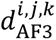 of inter-chain residue pairs from the AF3-predicted structures to construct the energy functions of the five corresponding populations. The energy function is defined as follows:

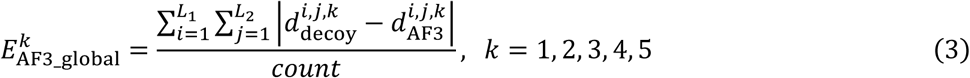

where *k* denotes that the population employs the distance map extracted from the *k-*th ranked model predicted by AF3.

Using the above-constructed energy functions as optimization targets, we designed a multi-population, multi-objective collaborative evolutionary algorithm based on the Monte Carlo sampling method^35^ (Details of the algorithmic workflow are described in **Supplementary Note 2**). Specifically, during the initialization phase, six independent populations were constructed using *E*_our_global_ and 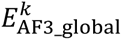, each population contains 100 solutions. These solutions are represented by a six-degree-of-freedom rigid-body transformation matrix, comprising three Euler angles for rotation and three spherical coordinates for translation, thereby describing a candidate protein conformation undergoing a relative spatial orientation change. For the six populations, the multi-objective optimization task is defined as follows:

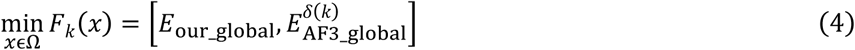

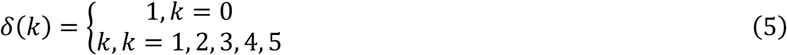

where *k* = 0 represents the multi-objective optimization task of the population which is constructed by our predicted distance maps. *k* = 1, 2, 3, 4, 5 representes the multi-objective optimization task of the *k*-th population which is constructed by AF3’s maps.

Subsequently, in the evolutionary phase, new candidate solutions were generated through evolutionary operators (mutation and crossover), leveraging the differences among individuals in the population to introduce random variations while enhancing population diversity. During the updating phase, each candidate solution was scored according to the objective energy function of its population and accepted into the solution set following the Metropolis criterion. The updated solution sets were then filtered using the Pareto dominance principle^36,37^, yielding a Pareto front that achieves cooperative optimization under multiple objective constraints. The Pareto front in this work is defined as follows:

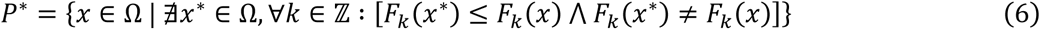

where, *x* denotes a candidate solution from the set of all possible solutions Ω. *x*^∗^represents a candidate solution to be compared with *x* for evaluating Pareto dominance. ℤ denotes the index set of objective functions, which in this work is {0, 1, 2, 3, 4, 5}. *F*_*k*_(*x*) represents the *k*-th objective energy function. For protein conformation optimization, each *x* ∈ Ω corresponds to a candidate conformation, and the *P*^∗^represents the Pareto front in the objective space, where each solution embodies a trade-off among multiple objectives, meaning that improvement in one objective inevitably leads to deterioration in at least one other. Details of the Pareto optimization principle is provided in the **Supplementary Note 3**.

Finally, the solutions filtered by the Pareto dominance principle were treated as an elite set and incorporated into a cyclic elitist replacement strategy, in which the elite solutions from population *k* replaced the same number of low-ranking individuals ranked by energy scores in population *k* + 1, while those from the last population replaced the low-ranking individuals in the first. The entire sampling process is performed for 300 iterations, and the Pareto elite sets obtained from each population in the final round are subjected to a confidence selection mechanism to identify the conformations with the lowest energy scores, which are then used as the inputs for the subsequent local exploitation stage.

### Local exploitation of CDRs based on local coordinate systems

Although the first stage has yielded antibody-antigen complex conformations with global conformational rationality and diverse spatial distributions, the high flexibility of CDRs remains a critical factor influencing interface recognition and binding affinity. Therefore, following the correction of relative orientations, the second stage focuses on local conformational optimization of the CDRs to enhance the physical plausibility and geometric complementarity of the interface conformations.

Based on the confidence selection mechanism using the energy function, the lowest-energy conformation from each population obtained after the complete first-stage process is used as the initial input. Using the pre-annotated start and end residue indexes of the antibody CDRs, we identified target regions for local perturbation. With the backbone dihedral angles of the residues as the optimization objects, the energy function is uniformly constructed using a residue-pair distance error function based on the local coordinate system. Specifically, for each residue in the antibody CDRs, a local orthogonal coordinate system is constructed with its C_β_ atom as the origin. The coordinates of the corresponding antigen residue in the sampled conformation are projected into this coordinate system and compared against the target distance constraints to quantify the spatial deviation of the antigen relative to the antibody’s local frame. The energy function is defined as follows:

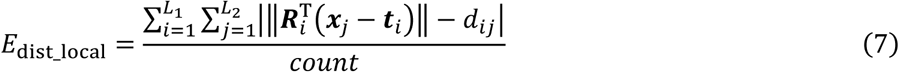

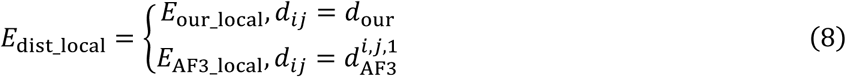

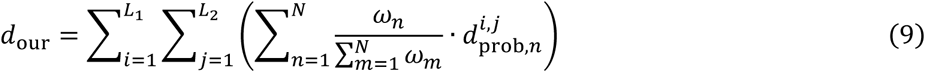

where 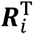 denotes the rotation matrix of the local orthogonal coordinate system constructed with antibody residue *i* as the reference; ***x***_*j*_ represents the C_β_ atom coordinates of antigen residue *j* in the sampled conformation; ***t***_*i*_ is the C_β_ atom coordinates of antibody residue *i*, serving as the origin of the local coordinate system; and *d*_*ij*_ is the target distance constraint taken from the inter-residue distance map generated by our predictor, in the same manner as in first Stage.

To further enhance the sensitivity and stability during the structural fine-tuning process, we constructed energy terms based on both the distance maps generated by our predictor and those derived from AF3’s top1 model, then integrated them through weighted fusion to form a unified local refinement energy function. The weighted energy function is defined as follows:

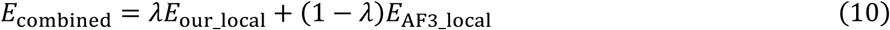

where the weighting coefficient *λ* is set to 0.5.

This weighted energy function is employed to constrain and evaluate the sampled conformations in the second stage, aiming to improve the physical plausibility and accuracy of the modeled structures. During the optimization process, Rosetta’s SmallMover protocol is applied to introduce large perturbations to the CDRs, generating multiple candidate conformations. Subsequently, each candidate conformation is applied to smaller perturbations, and a greedy selection strategy is adopted to retain lower-scoring conformations constrained by the weighted energy function *E*_combined_ in each iteration, thereby progressively refining the fine-grained geometry of the CDRs. This strategy enables efficient optimization of the CDR structures while maintaining the overall conformational stability. The algorithmic flow and detailed description of the second stage are provided in **Supplementary Note 4**.

### Energy-based conformation confidence selection mechanism

During the first-stage conformation sampling, each population independently evaluates its conformations based on the corresponding energy function. Specifically, for the population sampled using the distance maps predicted by DeepAAAssembly as the energy function, conformation quality is scored by *E*_our_global_. For the *k* -th population (*k* = 1, 2, 3, 4, 5) sampled using the distance maps derived from AF3, the corresponding energy function *E*^*k*^ is applied for scoring. Ultimately, the conformation with the lowest energy score from each population is selected as the initial input for the second-stage conformation sampling.

During the second-stage local refinement process, the weighted energy function *E*_combined_ is used to uniformly evaluate all candidate conformations. Upon completion of all iterations, the top-5 conformations with the lowest energy scores are selected as the final output.

### Evaluation metrics

We used two metrics to assess the quality of antibody-antigen complex models generated by DeepAAAssembly. The DockQ^43^ score evaluates the accuracy of the modeled interface regions and was calculated with the DockQ tool (https://github.com/bjornwallner/DockQ/). The TM-score^44^ measures global structural similarity between the generated model and the corresponding experimental structure and was calculated with the TM-score tool (https://zhanggroup.org/TM-score/). In addition to the above structural evaluation metrics, we further introduced the MAE to assess the performance of inter-chain residue distance prediction, which quantifies the average absolute deviation between the predicted and true inter-residue distances. Detailed definitions of these evaluation metrics are provided in **Supplementary Note 5**.

## Data availability

The authors declare that the data supporting the results and conclusions of this study are available within the paper and its Supplementary Information. The antibody-antigen data from DB5.5 dataset is available at: https://github.com/piercelab/antibody_benchmark. The CASP15-16 data including the experimental structures are available at: https://predictioncenter.org/casp15/index.cgi, https://predictioncenter.org/casp16/index.cgi.

## Code availability

DeepAAAssembly is available for download via GitHub at https://github.com/iobio-zjut/DeepAAAssembly.

## Acknowledgements

We thank members of the Guijun Zhang lab for discussion and feedback. Computational resources were provided by the College of Information Engineering at Zhejiang University of Technology. This work was supported by the National Key R&D Program of China [2022ZD0115103, G.Z.], the National Nature Science Foundation of China [62173304, G.Z.; 62573386, G.Z.], the “Pioneer” and “Leading Goose” R&D Program of Zhejiang [2025C01190, G.Z.], and the Zhejiang Province High-level Talent Special Support Program [2023R5248, G.Z.].

## Author contributions

G.Z. conceived and supervised the research. G.Z., S.W., and J.Z. designed the experiment. S.W., J.Z., X.C., Z.L., and D.H. collected the data and performed the experiment. G.Z., S.W., and J.Z. analyzed the data. S.W. and J.Z. wrote the manuscript, and G.Z. revised it. All authors read and approved the final manuscript.

## Additional information

Correspondence and requests for materials should be addressed to G.Z..

## Supplementary Information for

### Supplementary Figures

**Supplementary Fig. 1.**
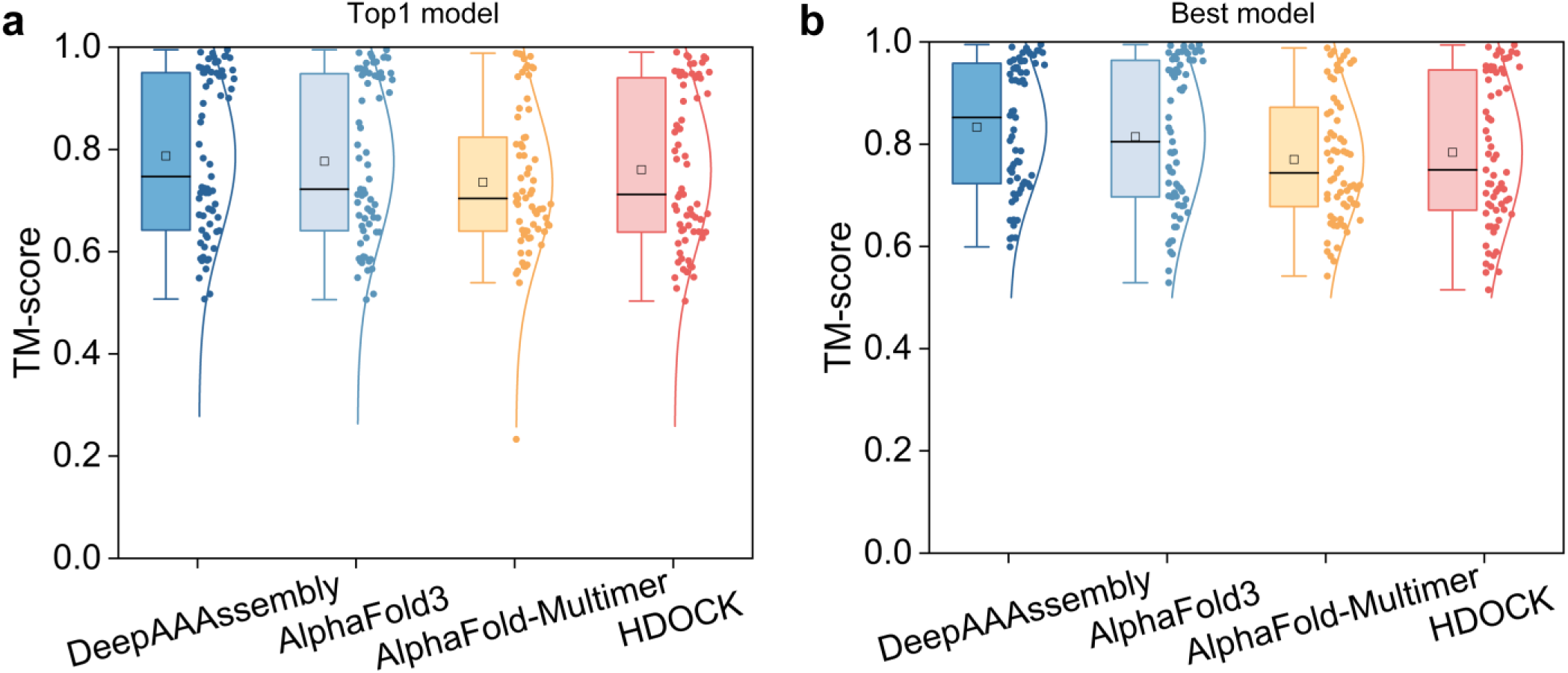
TM-score comparison among different methods. Comparison of the average TM-scores of top1 models (a) and best models (b) generated by DeepAAAssembly, AlphaFold3^1^, AlphaFold-Multimer^2^, and HDOCK^3^.

**Supplementary Fig. 2.**
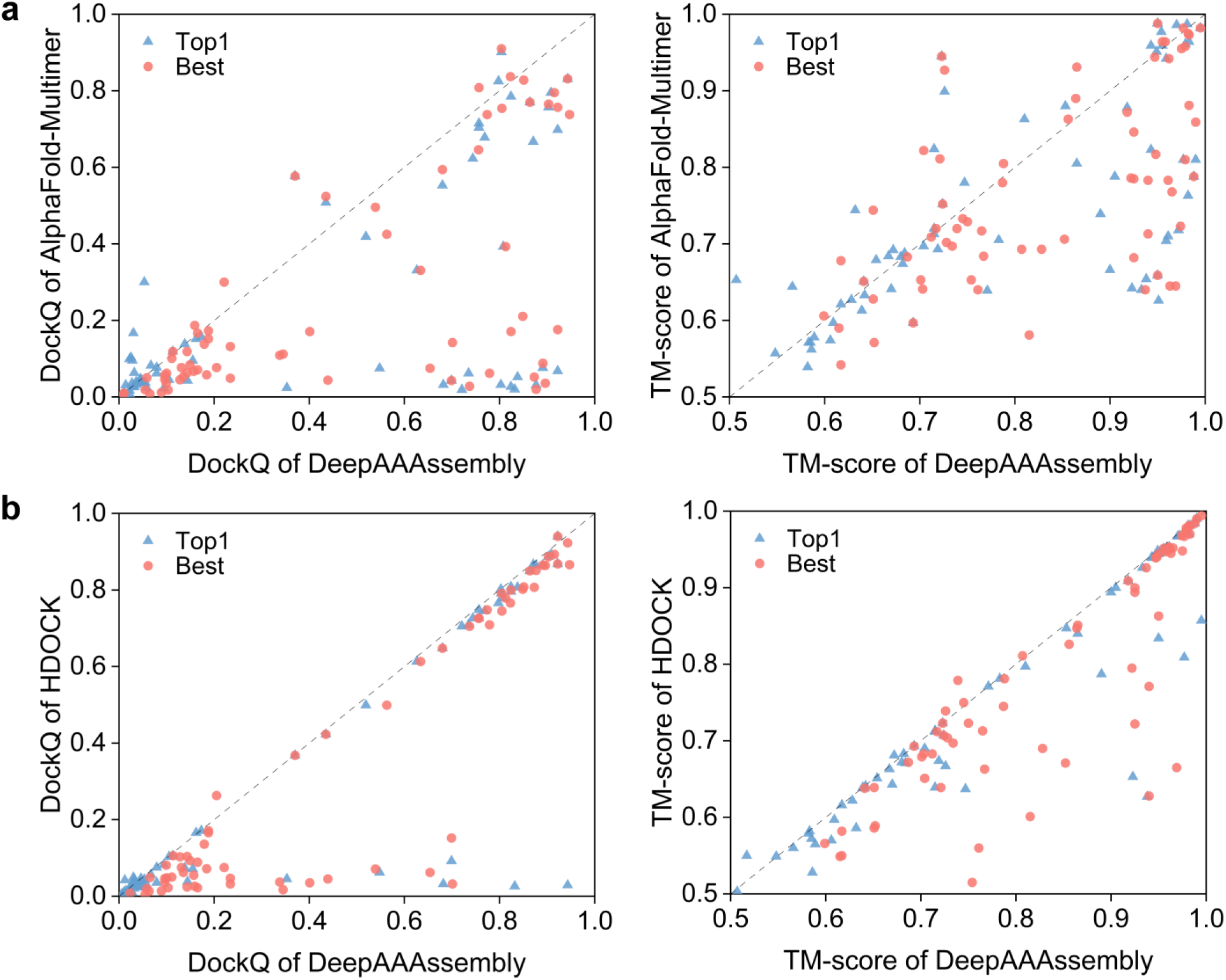
Performance comparison between DeepAAAssembly and the baseline methods AlphaFold-Multimer and HDOCK. **a,** DockQ and TM-score performance comparison between DeepAAAssembly and AlphaFold-Multimer across all test targets from the DB5.5^4^ benchmark set. **b,** DockQ and TM-score performance comparison between DeepAAAssembly and HDOCK across all test targets from the DB5.5 benchmark set.

**Supplementary Fig. 3.**
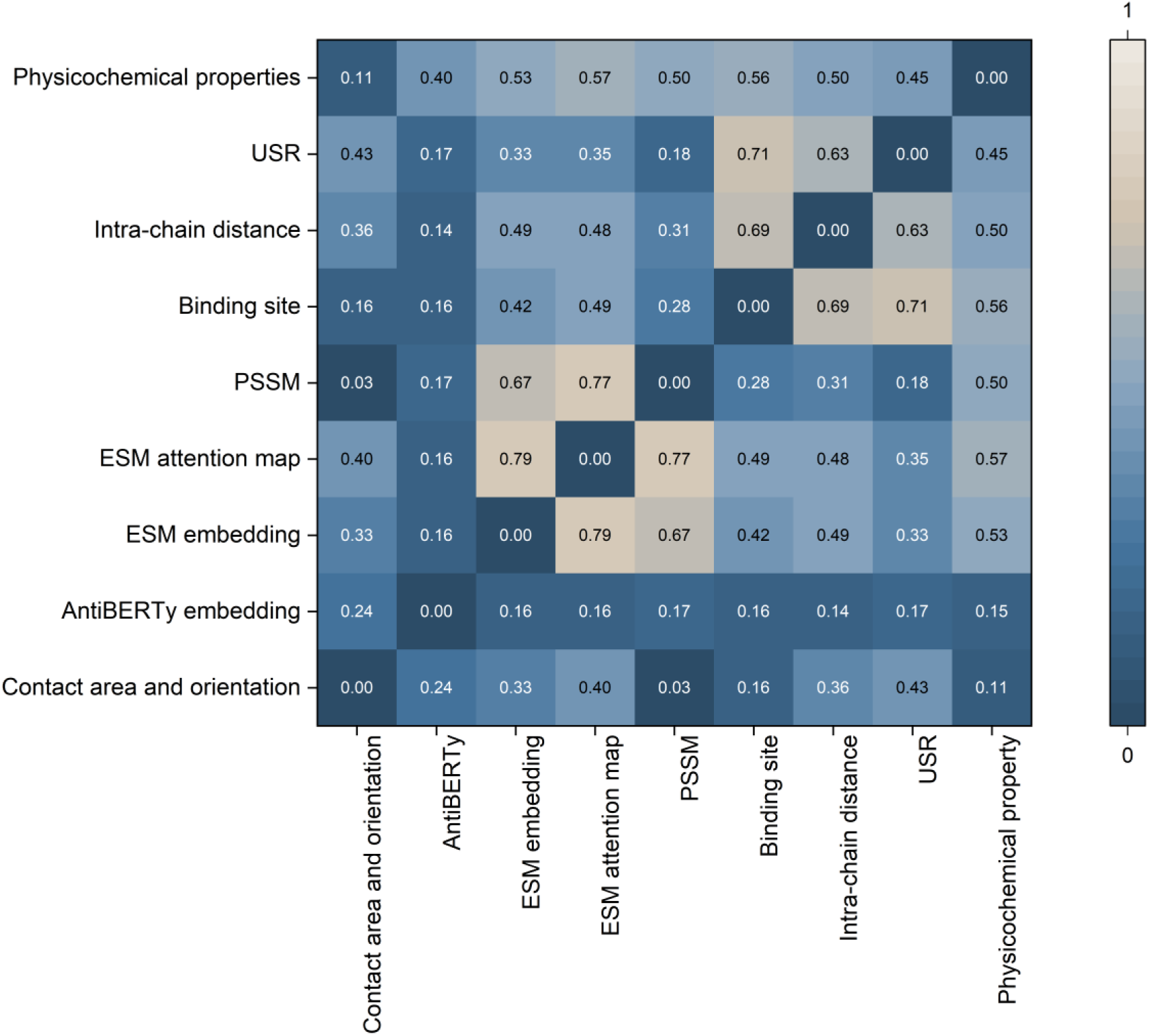
Correlation results of all features. All features of the antibody sequence were unified into a shape of [L, 1] through average pooling, and their correlations were computed using the Pearson correlation coefficient. The horizontal and vertical axes represent the feature types, where values closer to 0 (blue) indicate lower correlation, and values closer to 1 (gray) indicate higher correlation.

### Supplementary Notes

#### Supplementary Note 1. Calculation process of the residue level contact area and orientation features

The contact area and orientation features include the inter-residue contact surface area, solvent-accessible surface area of residues, and the directional characteristics of the inter-residue contact surfaces. We constructed residue-level contact area and orientation features based on atomic-level contact information to capture the interaction patterns between residues in antibody-antigen complexes. The schematic diagram is shown in **Supplementary Fig. 4**.

**Supplementary Fig. 4.**
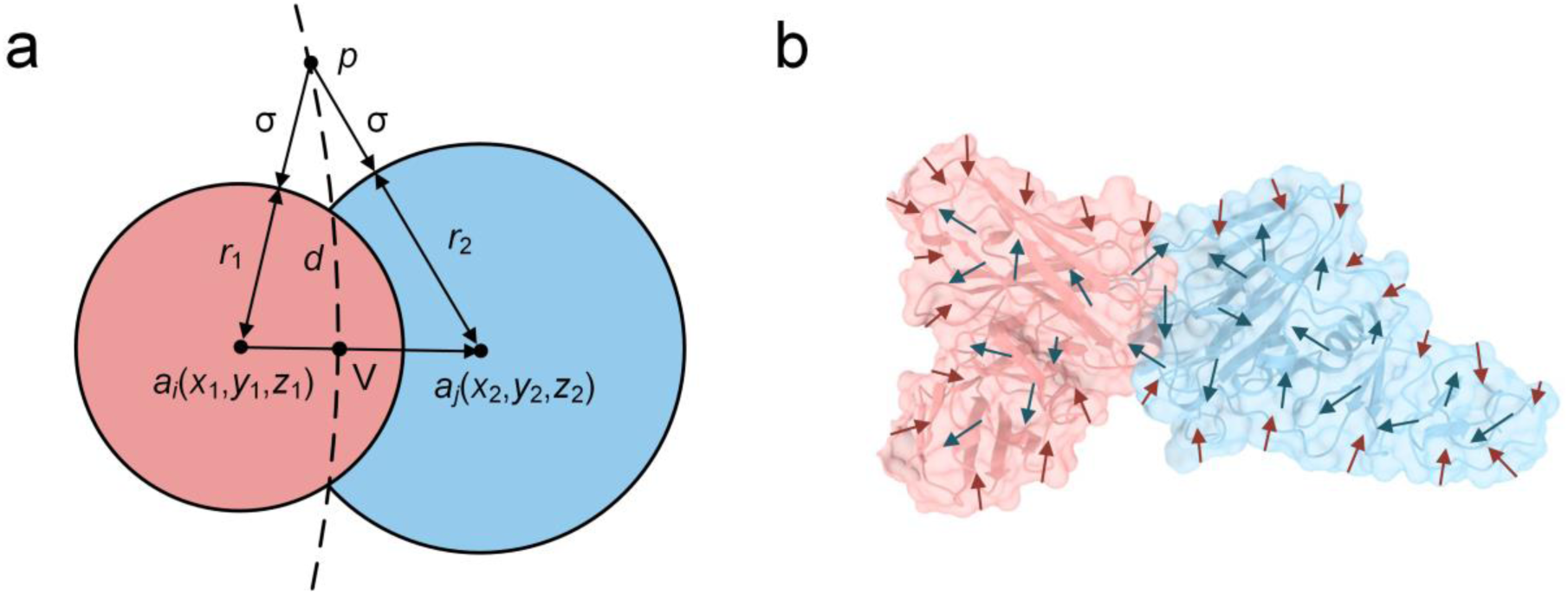
**a,** Schematic illustration of the contact between atoms *a*_*i*_ (red) and *a*_*j*_ (blue). *r*_1_and *r*_2_are the van der Waals radii of atoms *a*_*i*_ and *a*_*j*_, respectively; *d* is the interatomic distance; *p* is a point on the Voronoi surface, where the distances from *p* to the surfaces of atoms *a*_*i*_ and *a*_*j*_ are both σ. **b,** Schematic representation of residue-level contact surface orientations in an antibody-antigen complex. Red arrows indicate the contact surface orientations of surface residues, while blue arrows indicate those of buried residues.

First, we modeled atomic contact surfaces using the Voronoi^5^ tessellation algorithm and calculated the contact area between each atom pair *a*_*i*_ and *a*_*j*_ (ACA) via triangulation using the Voronota software (version 1.27.3834). Subsequently, the contact areas of all atom pairs belonging to the same residue pair *r*_*a*_ and *r*_*b*_ were summed to obtain the residue-level contact area (RCA):

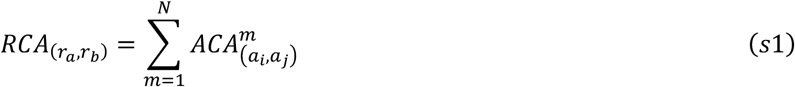

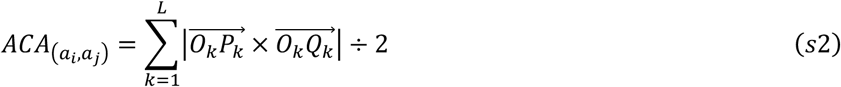

where, 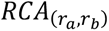 denotes the contact area between residues *r*_*a*_ and *r*_*b*_, 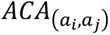 represents the contact surface area between atoms *a*_*i*_ and *a*_*j*_, *m* refers to the *m*-th atomic-level contact area, and *N* is the total number of atomic-level contacts between residues *r*_*a*_ and *r*_*b*_; *O*_*k*_, *P*_*k*_, *Q*_*k*_ corresponds to the vertices of the *k*-th triangle on the contact surface obtained via triangulation, and *L* is the total number of triangles on the contact surface. The residue-solvent contact area feature is obtained by summing the solvent-accessible areas of the constituent atoms.

We further introduced contact surface orientation vectors to characterize the spatial orientation of residues. For each pair of contacting atoms, we extract the support vector 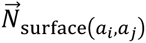 of their contact surface. The residue-level contact orientation vector is defined as the sum of all support vectors from the corresponding atom pairs:

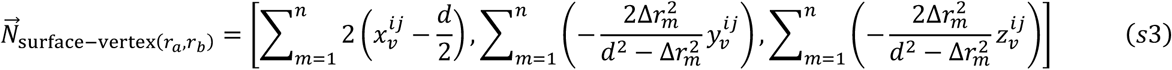

where, the coordinates of the reference point V are 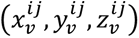, obtained at the intersection between the contact surface of the atom pair and the line connecting the two atoms; *d* represents the distance between the atom pair, *n* is the total number of atomic-level contacts between residues, and Δ*r*_*m*_ denotes the van der Waals radius difference of the *m*-th atomic-level contact surface.

#### Supplementary Note 2. Algorithm design of the first exploration stage

In the exploration-stage conformation sampling, we designed a multi-population, multi-objective cooperative optimization algorithm based on a differential evolution^6^ framework to comprehensively explore the possible relative spatial orientations of antibody-antigen complexes. The schematic flowchart of the exploration stage is shown in **Supplementary Fig. 5**.

**Supplementary Fig. 5.**
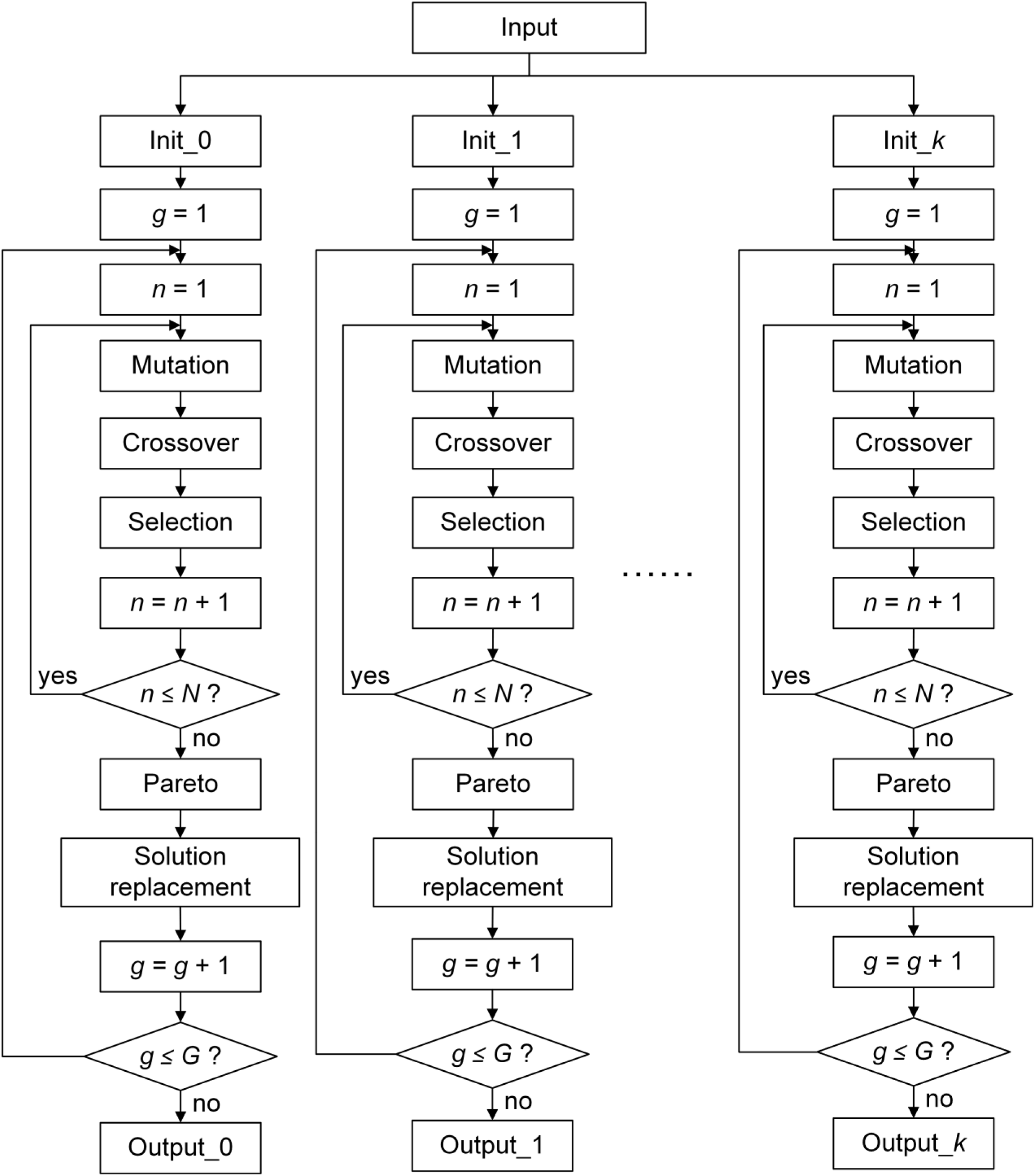
Schematic flowchart of the exploration stage.

We used the top1 model predicted by AF3 as the initial input and initialized six distinct populations by randomly generating *N* (*N* = 100) sets of Euler angles and spherical-coordinate translation parameters (six degrees of freedom in total). Each solution within a population represents a possible relative spatial conformation of the antibody-antigen complex. The conformation sampling of each population is guided by a specific energy function: one population’s energy function is constructed based on the inter-chain residue distance maps predicted by DeepAAAssembly, while the remaining five populations use energy functions derived from distance maps extracted from the five AF3-predicted structures, respectively. Each population independently undergoes the differential evolution process, with each iteration consisting of mutation, crossover, selection, and solution replacement, for a total of *G* (*G* = 300) iterations.

During the mutation stage, three distinct solutions are randomly selected from the initial population, and a mutant solution is generated by performing the following linear combination on the six parameters corresponding respectively to these three solutions:

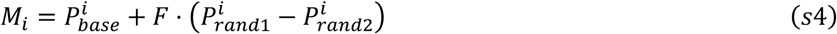

where, *M* represents the mutant solution; *i* denotes the *i* -th degree-of-freedom parameter; *P* indicates the original population; *base*, *rand*1, *rand*2 ∈ [1, *N*), and are three distinct indices corresponding to randomly selected solutions from *P*; and *F* = 0.5 is the scaling factor.

During the crossover stage, whether the mutant solution is retained is determined by the following condition:

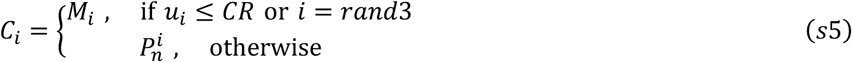

where, *C* represents the crossover solution; *u*_*i*_ denotes a uniformly distributed random number within the interval [0, 1); *CR* = 0.5 is the crossover rate; *rand*3 ∈ [1,6] indicates a randomly generated integer. This condition determines whether each of the six degrees-of-freedom parameters from the mutant solution *M* is accepted based on the crossover rate *CR*, while *rand*3 ensures that at least one parameter originates from the mutant solution. Otherwise, the corresponding parameter from the original solution *P*_*n*_ is retained.

In the selection stage, the Metropolis criterion is employed to determine whether the crossover solution is accepted into the next-generation population:

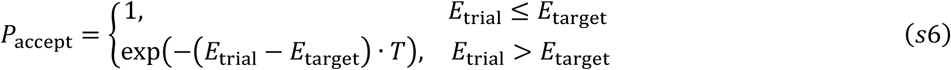

where, *E*_target_ denotes the energy score of the current solution, computed using either the energy function *E*_our_global_ or 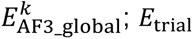 represents the energy score of the candidate crossover solution; and *T* is the thermal energy scale. If the candidate solution yields a lower energy, it is directly accepted; otherwise, it is accepted with a certain probability according to the Metropolis criterion.

During the solution replacement stage, a cyclic elitist replacement strategy was applied, where the elite conformations (Pareto front) obtained after multi-objective optimization in population *k* replaced an equal number of low-ranking conformations (ranked by energy scores) in population *k* + 1, and the elite conformations from the last population replaced the low-ranking conformations in the first. In the multi-objective optimization process, the algorithm simultaneously optimized multiple energy functions and employed the Pareto dominance principle to retain conformations not dominated by others while filtering out inferior ones, thereby guiding the evolutionary direction of the populations. After the final iteration, the Pareto fronts obtained from each population were evaluated using the energy-based confidence selection mechanism, and the lowest-energy conformation from each population was selected as the input for the subsequent local exploitation stage. The detailed description of the Pareto dominance principle is shown in the **Supplementary Note 3**.

#### Supplementary Note 3. Description of the Pareto dominance principle

**Supplementary Fig. 6** illustrates the geometric representation of Pareto dominance principle^7,8^. The horizontal and vertical axes correspond to two objective functions, *E*_1_and *E*_2_, respectively. Point *x*_0_ denotes the initial solution, whose objective function values are (e_1_, e_2_). Taking *x*_0_ as the origin, the plane is divided into four quadrants to analyze the dominance relationships among candidate solutions under different combinations of objective values. For any two solutions *x*_*i*_ and *x*_*j*_, if *E*_*k*_(*x*_*i*_) ≤ *E*_*k*_(*x*_*j*_) for all *k*, and there exists at least one *k* such that *E*_*k*_(*x*_*i*_) < *E*_*k*_(*x*_*j*_), then *x*_*i*_ is said to dominate *x*_*j*_. Conversely, if the two solutions perform differently across objectives, with each being better in at least one objective, they are considered non-dominated with respect to each other.

**Supplementary Fig. 6.**
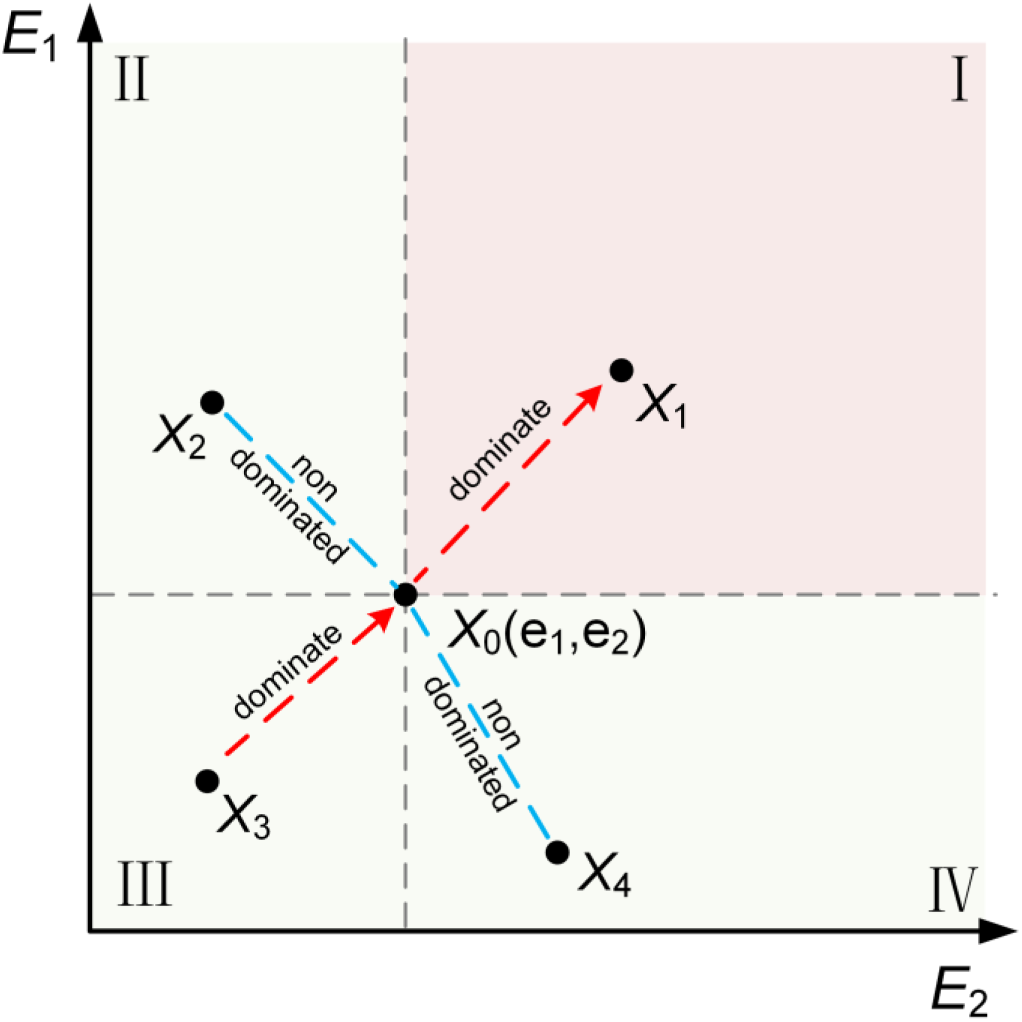
Schematic illustration of the Pareto front. The horizontal and vertical axes represent two objective functions, *E*_1_ and *E*_2_, respectively. Point *x*_0_ denotes the initial solution with corresponding objective values (e_1_, e_2_). Taking *x*_0_as the origin, the plane is divided into four quadrants. The red region (first quadrant) represents solutions that are inferior to *x*_0_in both objectives—these are dominated by *x*_0_ and are therefore excluded from the Pareto front. The green regions (second, third, and fourth quadrants) correspond to solutions that are better than *x*_0_ in at least one objective—these are non-dominated with respect to *x*_0_ and can be accepted into the Pareto front. In the figure, red dashed arrows indicate the direction of dominance, pointing from the dominating solution to the dominated one, while blue dashed lines denote non-dominated pairs of solutions.

According to this definition, the first quadrant (red-shaded area) represents solutions that are inferior to *x*_0_in all objectives; these candidates are dominated by *x*_0_ and are thus excluded from the Pareto front. The second, third, and fourth quadrants (green-shaded areas) correspond to cases where the candidate solution outperforms *x*_0_ in at least one objective. Therefore, these solutions are non-dominated with respect to *x*_0_and are accepted into the Pareto front. For example, if candidate *x*_1_ lies in the first quadrant, it performs no better than *x*_0_ in either *E*_1_ or *E*_2_, hence *x*_0_ dominates *x*_1_, and *x*_1_ is excluded from the Pareto front. If *x*_2_ lies in the second quadrant where *E*_1_(*x*_2_) > *E*_1_(*x*_0_) and *E*_2_(*x*_2_) < *E*_2_(*x*_0_), the two are non-dominated, so *x*_2_ is accepted. Symmetrically, a solution *x*_4_ in the fourth quadrant is also non-dominated with respect to *x*_0_, and thus accepted. If *x*_3_falls in the third quadrant where *E*_1_(*x*_3_) < *E*_1_(*x*_0_) and *E*_2_(*x*_3_) < *E*_2_(*x*_0_), then *x*_3_ dominates *x*_0_, *x*_3_ is accepted, and *x*_0_ must be removed from the Pareto front. Before any newly generated candidate is added to the Pareto front, its dominance relationship with all existing solutions must be evaluated to determine acceptance. As shown by the green-shaded regions, every accepted solution achieves improvement in at least one objective, ensuring that the multi-objective optimization process remains non-degrading and cooperatively improving in the Pareto dominance principle.

#### Supplementary Note 4. Algorithm design of the second exploitation stage

In the second-stage conformation sampling, we designed a two-step local refinement strategy based on the SmallMover protocol in the Rosetta^9^ software (see **Supplementary Fig. 7** for a schematic illustration). This strategy consists of a large-amplitude perturbation stage followed by a fine-tuning stage, enabling a progressive transition from local search to precise optimization while maintaining physical plausibility. The SmallMover protocol perturbs backbone dihedral angles within a defined small range to achieve local conformation sampling. Its key advantage lies in the use of a “movemap” object, which precisely restricts the perturbation region, thereby effectively preventing structural deviations from propagating into unrelated parts of the protein. The key parameters in the SmallMover protocol include *kT*, which specifies the Monte Carlo (MC) acceptance temperature (evaluated only using the Rama score, thus not a full MC simulation); *n_moves*, which defines the number of consecutive moves to perform; and *angle_max*, which determines the maximum backbone perturbation angle.

**Supplementary Fig. 7.**
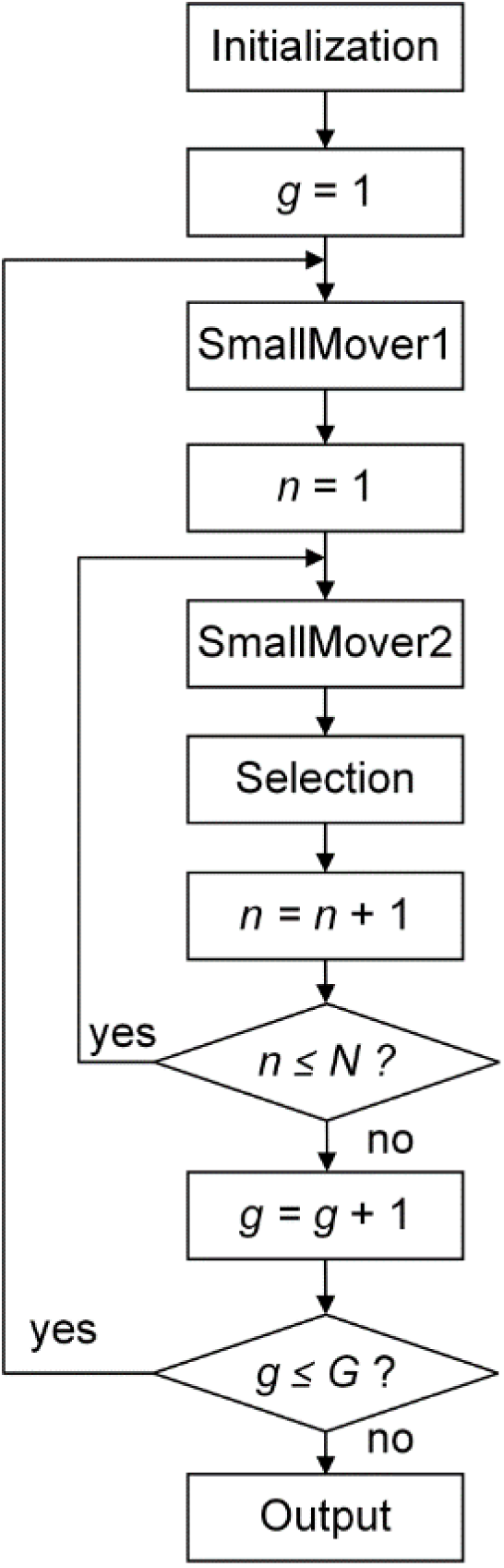
Schematic flowchart of the exploitation stage. In this stage, the optimal conformations obtained from the first-stage global sampling were used as inputs, and each conformation was iteratively optimized over *G* rounds (*G* = 20). The movemap settings were defined based on the complementarity-determining region (CDR) indices identified by ANARCI^10^ under the IMGT^11^ numbering scheme.

During the large-amplitude perturbation stage, conformations were evaluated using the default *rama*^3^ score to ensure physical plausibility, and accepted conformations were added to the population. The specific parameters were set to an acceptance temperature of *kT* = 5, a maximum torsion angle of *angle_max* = 10°, and *n_moves* = 5 continuous perturbations per iteration, promoting broad exploration of the conformational space. Subsequently, in the fine-adjustment stage, the scoring function was switched to the distance-constrained energy function *E*_combined_, which integrates our model-predicted distance maps with those derived from AF3 as directional guidance for perturbation. The parameters were adjusted to *kT* = 2, *angle_max* = 2°, and *n_moves* = 1 to enable subtle, localized optimization. After each perturbation, *E*_combined_ was used to evaluate whether the conformation update should be accepted. Finally, after *G* rounds of optimization for each input structure, the resulting conformations were ranked based on their *E*_combined_ energy scores, and the best-performing conformations were selected as outputs.

#### Supplementary Note 5. Details of evaluation metrics

##### DockQ

DockQ^12^ is a metric that integrates *F*_nat_, *LRMS*, and *iRMS* into a single continuous score ranging from 0 to 1, designed to quantify the quality of protein complex structural models. Higher values indicate greater agreement between the predicted and native structures. The calculation formula is as follows:

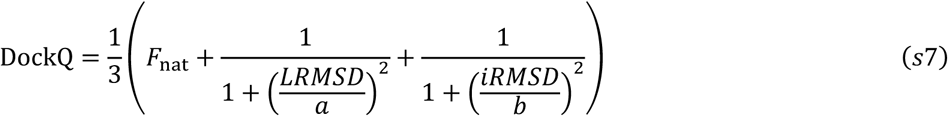

where, *F*_nat_ represents the fraction of interface residues in the predicted structure that overlap with those in the native structure; *LRMSD* represents the root-mean-square deviation of the ligand backbone (the shorter chain in the complex) after superimposing it onto the longer chain (receptor); *iRMSD* is the root-mean-square deviation of the interface backbone residues. The scaling factors for *LRMSD* and *iRMSD* are *a* = 8.5 and *b* = 1.5, respectively.

##### TM-score

TM-score^13^ is a metric used to evaluate the topological similarity between protein structures, ranging from 0 to 1, with higher values indicating greater structural accuracy. The calculation formula is as follows:

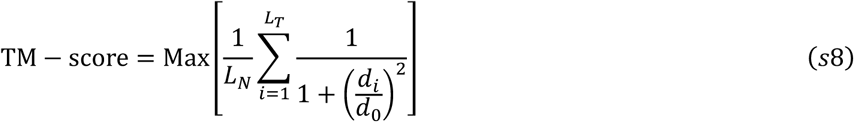

where, *L*_*N*_ is the length of the native structure, *L*_*T*_ is the number of residues aligned with the template structure, *d*_*i*_ is the distance between the *i*-th pair of aligned residues, and *d*_0_ is a scaling parameter used to normalize the deviation; “Max” indicates taking the maximum value after optimal spatial superposition.

##### MAE

The mean absolute error represents the average of the absolute differences between the true distances and the predicted distances, and is calculated as follows:

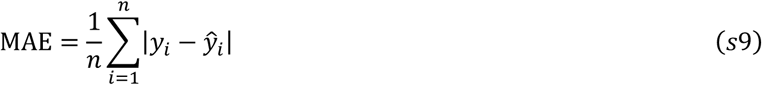

where, *y*_*i*_ denotes the true value, *y*^_*i*_ denotes the predicted value, and *n* is the number of samples.

### Supplementary Tables

**Supplementary Table 1.** Detailed description of input features for the network.

| Category | Feature & Shape | Description |
| --- | --- | --- |
| **Physicochemical features** | Contact area and orientation<br>[L, 5] | Residue-residue contact areas and residue solvent-accessible surface areas are calculated using the Voronoi tessellation model of atomic interfaces, followed by triangulation analysis implemented in the Voronota<sup>5</sup> software. For each residue, a contact orientation vector is computed as the vector sum of the directions toward all neighboring contact residues, representing the overall spatial orientation of local interactions. |
|  | Physical and chemical property<br>[L, 7] | The physical and chemical properties reflect the intrinsic characteristics of each amino acid and consist of the following seven attributes<sup>14</sup>: Steric parameter, Polarity, Volume, Hydrophobicity, Isoelectric point, Helix probability, and Sheet probability. |
| **Evolutionary features** | ESM attention map<br>[L, L, 144] | The attention maps are generated by the protein language model ESM-MSA-1b<sup>15</sup> to represent the potential spatial proximity between inter-chain residues and the relationships among functionally critical residues. |
|  | ESM embedding<br>[L, 768] | The ESM embeddings, produced by ESM-MSA-1b from the MSA, encode contextual information and capture evolutionary and conservation patterns of amino acid sequences, as well as potential interaction features. |
|  | AntiBERTy embedding<br>[L, 512] | Generated by the BERT-based pretrained language model AntiBERTy<sup>16</sup>, which is specifically trained on antibody sequences, these embeddings are used to capture the distinctive features of antibodies. |
|  | Position-specific scoring matrix<br>[L, 20] | The position-specific scoring matrix<sup>17</sup> (PSSM) reflects the evolutionary conservation and mutation propensity of amino acid sequences, indirectly revealing the importance of residues, their functional tendencies, and potential interaction features. |
| **Geometric features** | Ultrafast shape recognition<br>[L, 3] | During the computation of Ultrafast Shape Recognition<sup>18</sup>, the $i$-th residue $r_i$ in a monomer is selected as the reference point. The residue $c_i$ that is farthest from $r_i$ in Euclidean space is then identified, followed by locating the residue $h_i$ that is farthest from $c_i$. This procedure characterizes both local and global topological relationships within the protein structure. |
|  | Intra-chain distance<br>[L, L, 64] | The intra-chain residue distance is derived from the distance map between different residues within a monomeric protein, which can to some extent reflect its folding pattern. |
|  | Binding site<br>[L, 1] | The binding site is generated by the deep learning model PeSTo<sup>19</sup>, which outputs the probability of each residue participating in interface formation, enabling the model to more precisely focus on key residues likely involved in the interaction interface. |

**Supplementary Table 2.**
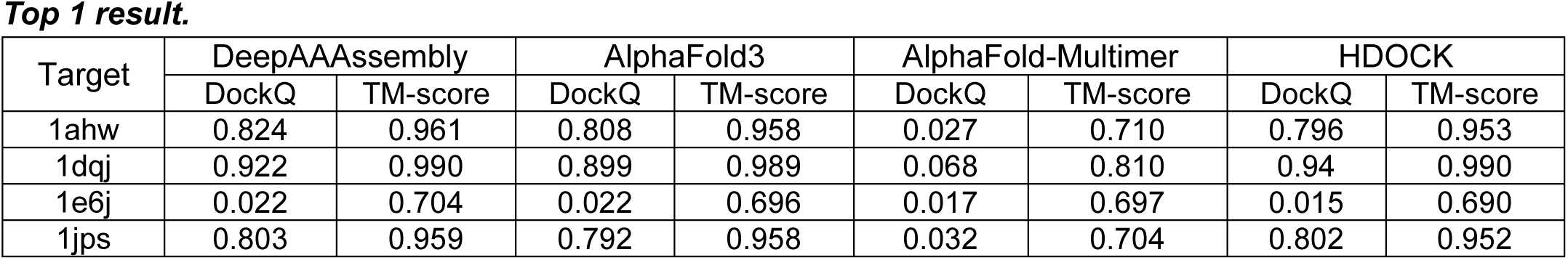
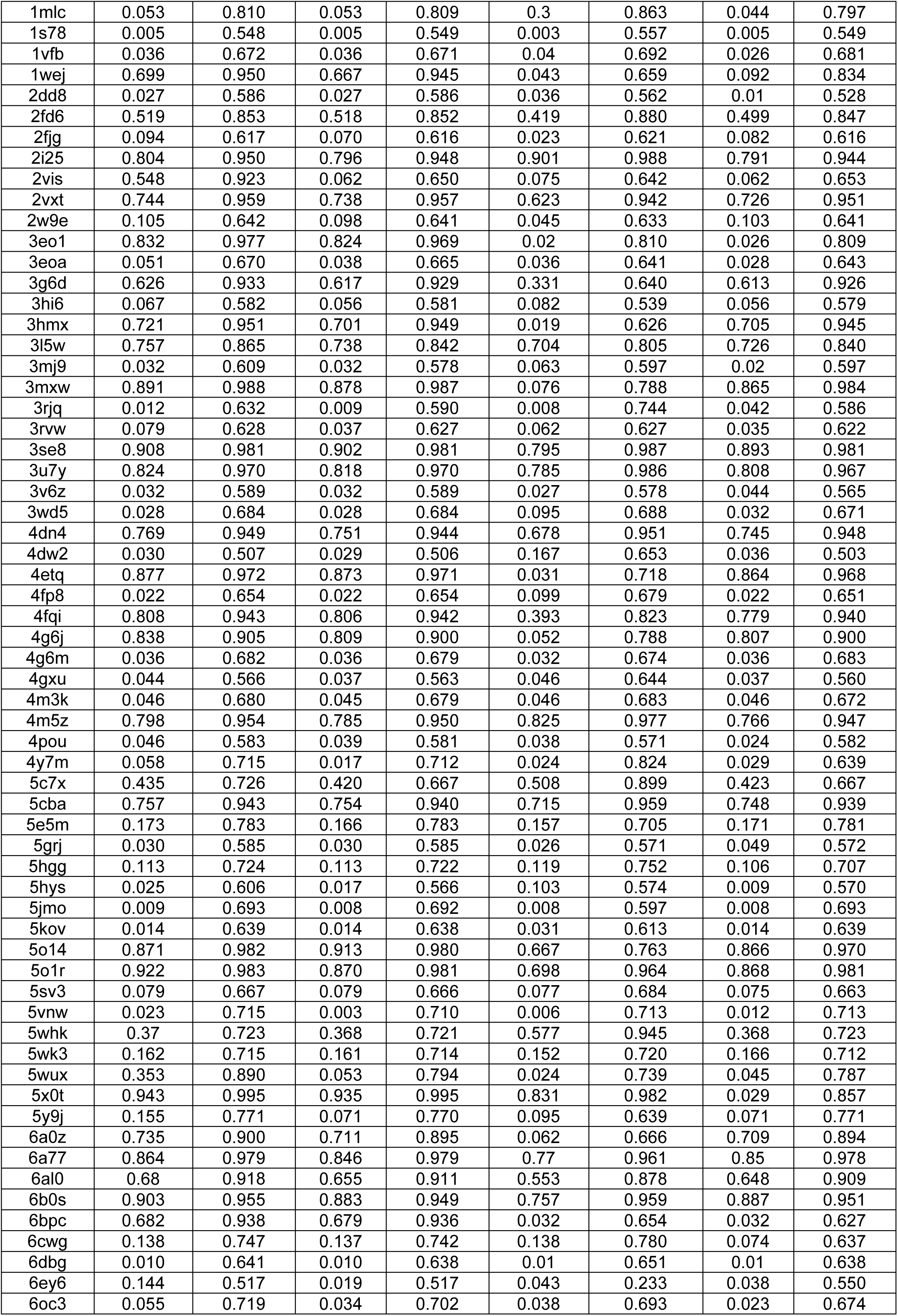
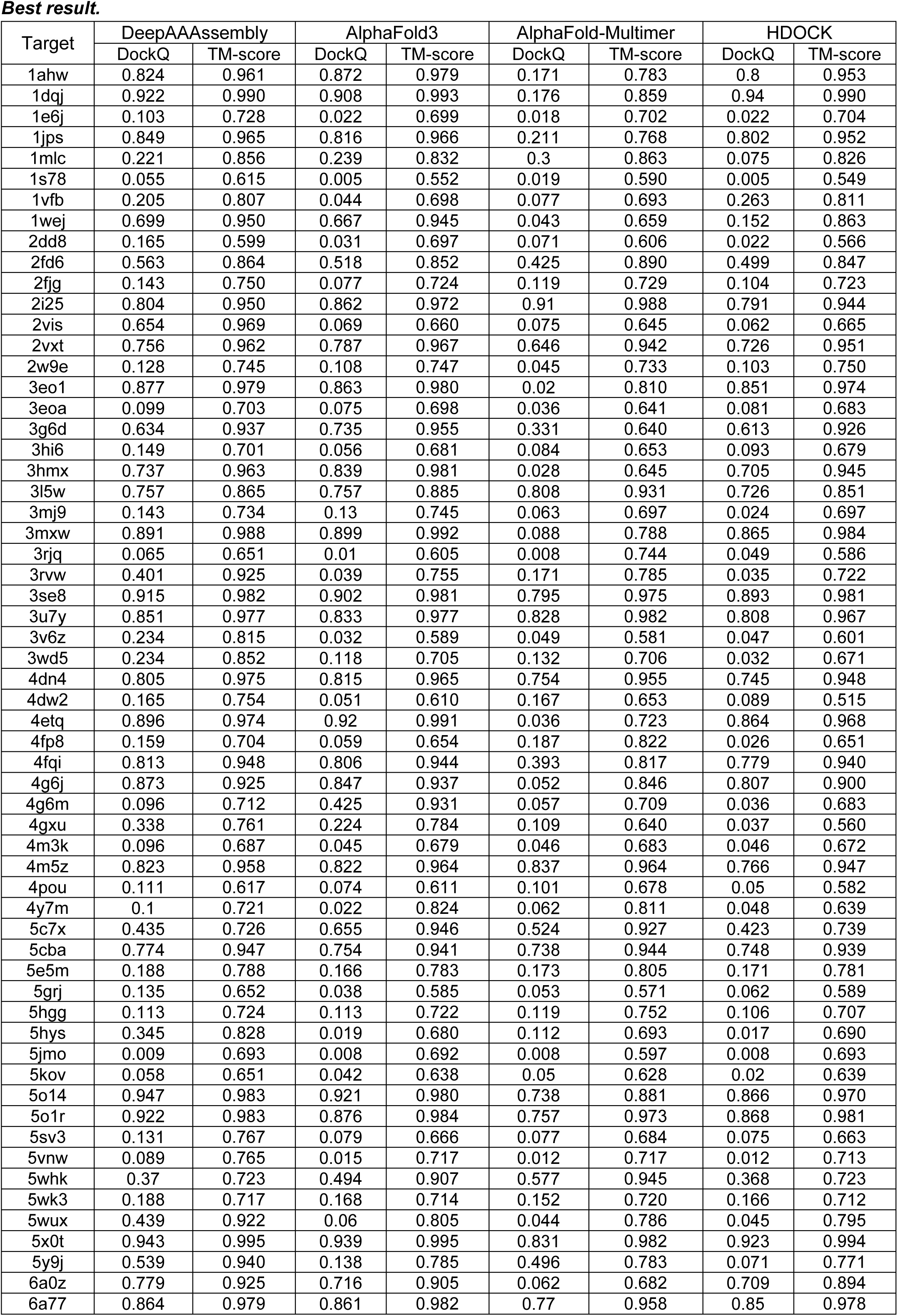
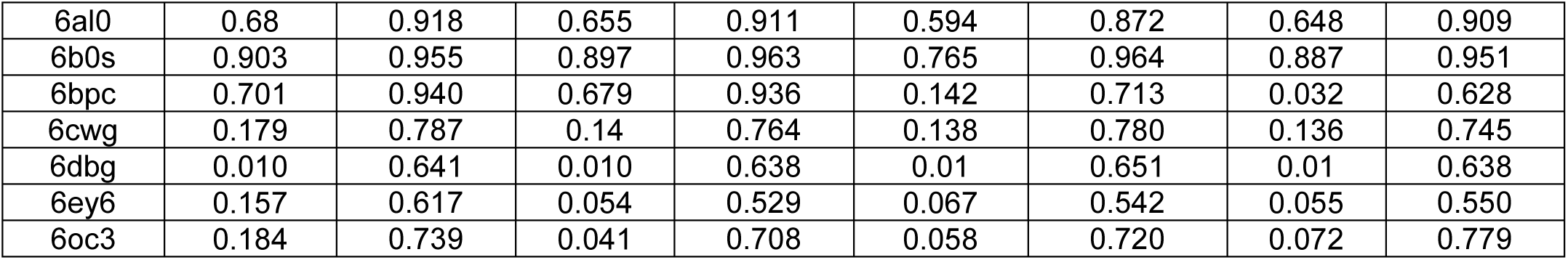
Specific test results of DB5.5 benchmark set.

**Supplementary Table 3.** HHblits^20^ Parameters.

| Parameter | Value | Description |
| --- | --- | --- |
| -cpu | 10 | Number of CPUs used |
| -n | 3 | Number of iterations run |
| -e | 0.001 | E-value threshold for result arrangement |
| -id | 99 | Maximum sequence consistency |
| -cov | 0.4 | Minimum coverage of the target sequence |

**Supplementary Table 4.** Weight setting corresponding to distance bin.

| Bin | Weight |
| --- | --- |
| $1 \leq c \leq 12$ | 1 |
| $13 \leq c \leq 20$ | 0.7 |
| $21 \leq c \leq 28$ | 0.5 |
| $c=0$ / $c \geq 29$ | 0.1 |

### Supplementary Algorithms

#### Supplementary Algorithm 1.

**Pseudocode of the Feature Preprocessing Module**

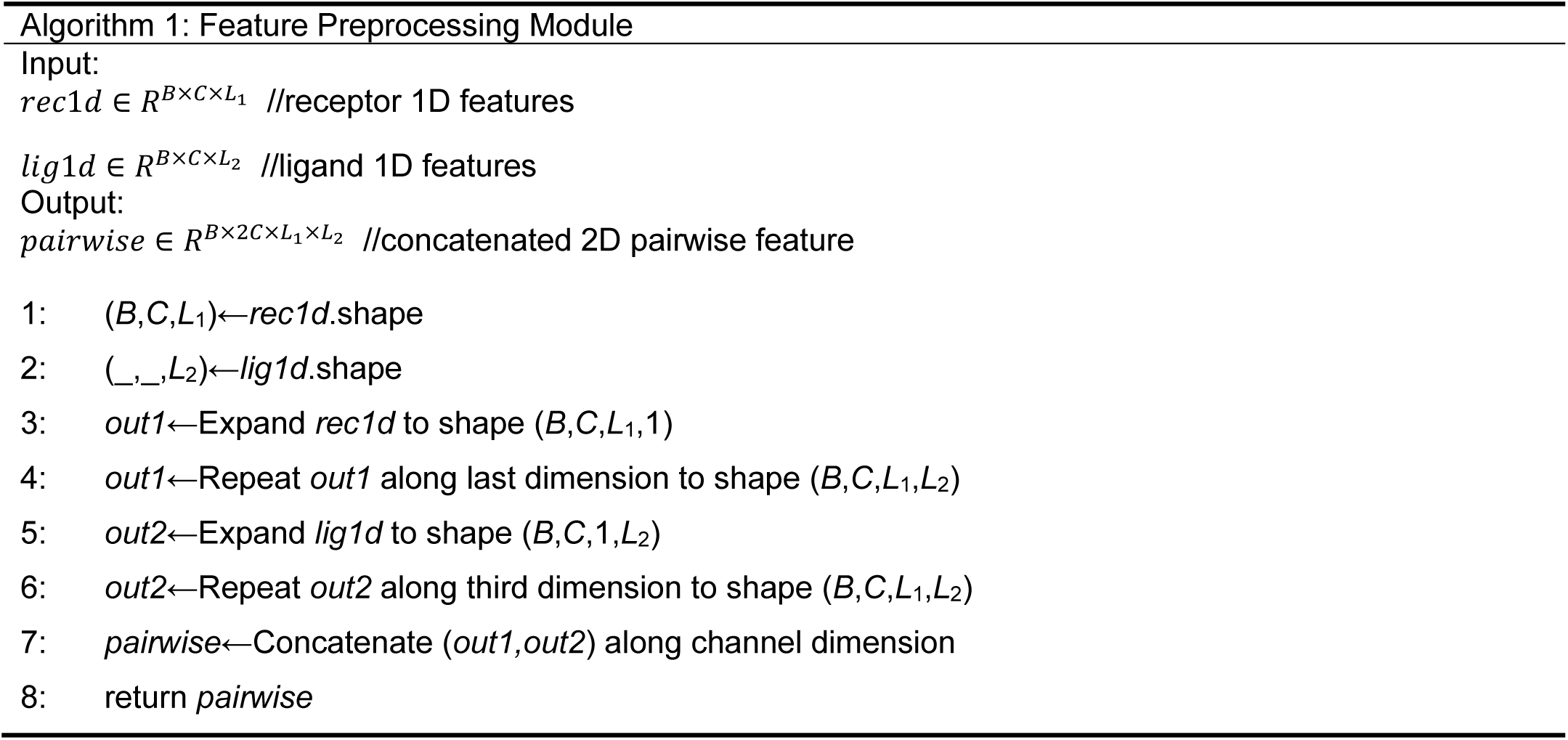

#### Supplementary Algorithm 2.

**Pseudocode of the Multi-Column Convolutional Neural Network Module**

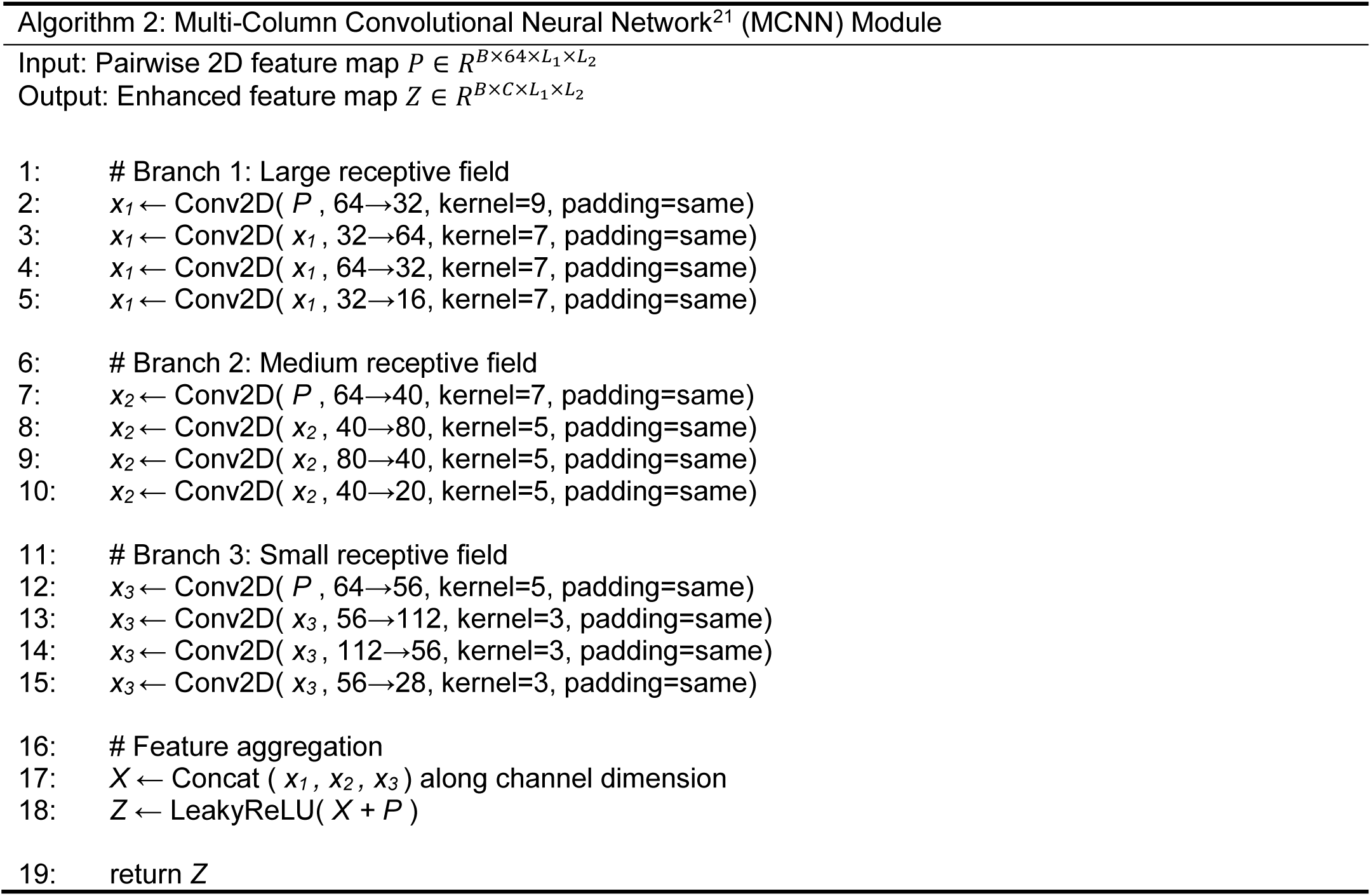

#### Supplementary Algorithm 3.

**Pseudocode for the Triangular multiplicative update layer**

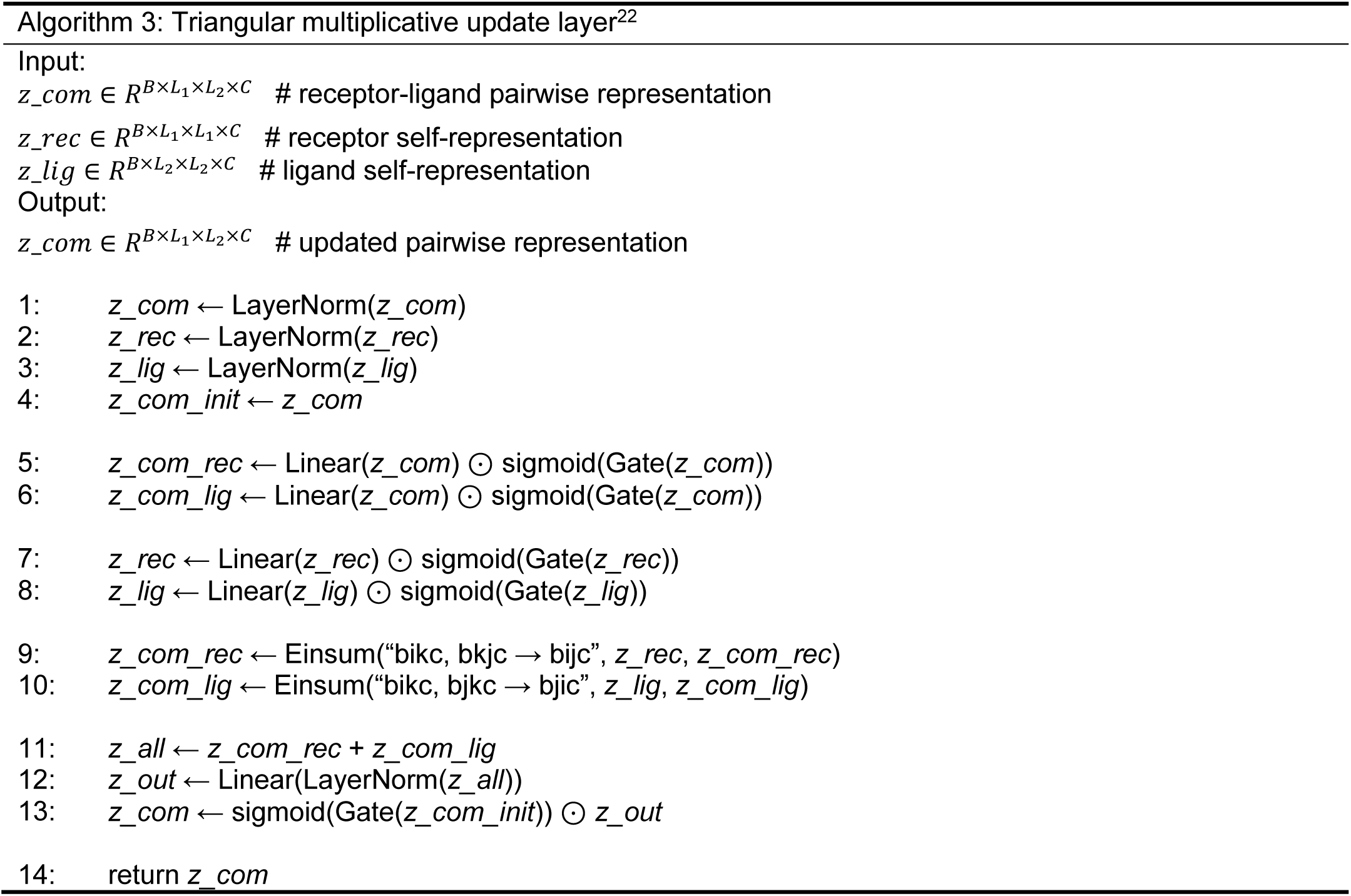

#### Supplementary Algorithm 4.

**Pseudocode of the Triangular self-attention layer**

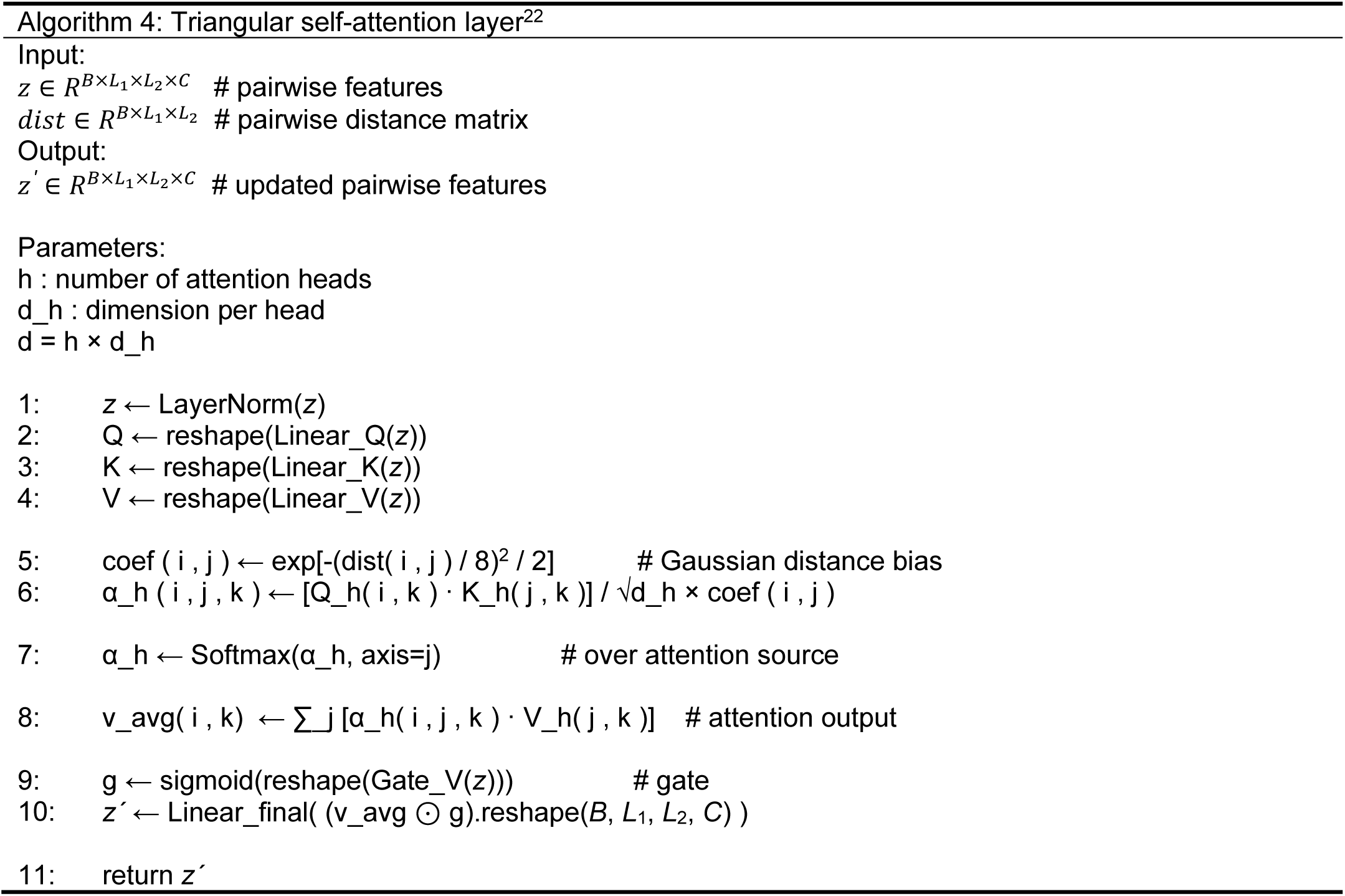

## References

1 Pantaleo, G., Correia, B., Fenwick, C., Joo, V. S. & Perez, L. Antibodies to combat viral infections: development strategies and progress. Nat. Rev. Drug Discov. 21, 676–696 (2022).

2 Passon, M. et al. Stability convergence in natural antibodies with ultra-long hypervariable loops. *Commun*. Biol. 8, 635 (2025).

3 Gaudreault, F., Corbeil, C. R. & Sulea, T. Enhanced antibody-antigen structure prediction from molecular docking using AlphaFold2. Sci. Rep. 13, 15107 (2023).

4 Yin, R., Feng, B. Y., Varshney, A. & Pierce, B. G. Benchmarking AlphaFold for protein complex modeling reveals accuracy determinants. Protein Sci. 31, e4379 (2022).

5 Jumper, J. et al. Highly accurate protein structure prediction with AlphaFold. Nature 596, 583–589 (2021).

6 Evans, R. et al. Protein complex prediction with AlphaFold-Multimer. Preprint at biorxiv, 10.1101/2021.10.04.463034 (2021).

7 Baek, M. et al. Accurate prediction of protein structures and interactions using a three-track neural network. Science 373, 871–876 (2021).

8 Lin, Z. et al. Evolutionary-scale prediction of atomic-level protein structure with a language model. Science 379, 1123–1130 (2023).

9 Mirdita, M. et al. ColabFold: making protein folding accessible to all. Nat. Methods 19, 679–682 (2022).

10 Zheng, W. et al. Improving deep learning protein monomer and complex structure prediction using DeepMSA2 with huge metagenomics data. Nat. Methods 21, 279–289 (2024).

11 Bryant, P., Kelkar, A., Guljas, A., Clementi, C. & Noé, F. Structure prediction of protein-ligand complexes from sequence information with Umol. Nat. Commun. 15, 4536 (2024).

12 Qiao, Z., Nie, W., Vahdat, A., Miller III, T. F. & Anandkumar, A. State-specific protein–ligand complex structure prediction with a multiscale deep generative model. *Nat*. Mach. Intell. 6, 195–208 (2024).

13 Abramson, J. et al. Accurate structure prediction of biomolecular interactions with AlphaFold 3. Nature 630, 493–500 (2024).

14 Gaudreault, F., Sulea, T. & Corbeil, C. R. AI-augmented physics-based docking for antibody-antigen complex prediction. Bioinformatics 41, btaf129 (2025).

15 Giulini, M. et al. Towards the accurate modelling of antibody− antigen complexes from sequence using machine learning and information-driven docking. Bioinformatics 40, btae583 (2024).

16 Stahl, K. et al. Modelling protein complexes with crosslinking mass spectrometry and deep learning. Nat. Commun. 15, 7866 (2024).

17 Feng, S. et al. Integrated structure prediction of protein–protein docking with experimental restraints using ColabDock. *Nat*. Mach. Intell. 6, 924–935 (2024).

18 McGuffin, L. J. et al. Prediction and quality assessment of protein quaternary structure models using the MultiFOLD2 and ModFOLDdock2 servers. Nucleic Acids Res. 53, W472–W477 (2025).

19 Raouraoua, N. et al. MassiveFold: unveiling AlphaFold’s hidden potential with optimized and parallelized massive sampling. Nat. Comput. Sci. 4, 824–828 (2024).

20 Chen, R., Li, L. & Weng, Z. ZDOCK: an initial-stage protein-docking algorithm. *Proteins Struct., Funct.*, Bioinf. 52, 80–87 (2003).

21 Gray, J. J. et al. Protein–protein docking with simultaneous optimization of rigid-body displacement and side-chain conformations. J. Mol. Biol. 331, 281–299 (2003).

22 Yan, Y., Tao, H., He, J. & Huang, S.-Y. The HDOCK server for integrated protein–protein docking. Nat. Protoc. 15, 1829–1852 (2020).

23 Sircar, A. & Gray, J. J. SnugDock: paratope structural optimization during antibody-antigen docking compensates for errors in antibody homology models. PLoS Comput. Biol. 6, e1000644 (2010).

24 Jiménez-García, B. et al. LightDock: a new multi-scale approach to protein–protein docking. Bioinformatics 34, 49–55 (2018).

25 Olechnovič, K. & Venclovas, Č. VoroIF-GNN: Voronoi tessellation-derived protein–protein interface assessment using a graph neural network. *Proteins Struct., Funct.*, Bioinf. 91, 1879–1888 (2023).

26 Gao, Z. et al. Hierarchical graph learning for protein–protein interaction. Nat. Commun. 14, 1093 (2023).

27 Rao, R. et al. MSA Transformer. Proceedings of the 38th International Conference on Machine Learning. 139, 8844–8856 (2021).

28 Ruffolo, J. A., Gray, J. J. & Sulam, J. Deciphering antibody affinity maturation with language models and weakly supervised learning. Preprint at arXiv, 10.48550/arXiv.2112.07782 (2021).

29 Altschul, S. F. et al. Gapped BLAST and PSI-BLAST: a new generation of protein database search programs. Nucleic Acids Res. 25, 3389–3402 (1997).

30 Ballester, P. J. & Richards, W. G. Ultrafast shape recognition to search compound databases for similar molecular shapes. J. Comput. Chem. 28, 1711–1723 (2007).

31 Krapp, L. F., Abriata, L. A., Cortés Rodriguez, F. & Dal Peraro, M. PeSTo: parameter-free geometric deep learning for accurate prediction of protein binding interfaces. Nat. Commun. 14, 2175 (2023).

32 Zhang, Y., Zhou, D., Chen, S., Gao, S. & Ma, Y. Single-image crowd counting via multi-column convolutional neural network. Proceedings of the IEEE conference on computer vision and pattern recognition, 589–597 (2016).

33 Lin, P., Tao, H., Li, H. & Huang, S.-Y. Protein–protein contact prediction by geometric triangle-aware protein language models. *Nat*. Mach. Intell. 5, 1275–1284 (2023).

34 Parzen, E. On estimation of a probability density function and mode. Ann. Math. Stat. 33, 1065–1076 (1962).

35 Metropolis, N., Rosenbluth, A. W., Rosenbluth, M. N., Teller, A. H. & Teller, E. Equation of state calculations by fast computing machines. J. Chem. Phys. 21, 1087–1092 (1953).

36 Meng, Z. et al. Application of state-of-the-art multiobjective metaheuristic algorithms in reliability-based design optimization: a comparative study. Struct. Multidiscip. Optim. 66, 191 (2023).

37 Panagant, N., Pholdee, N., Bureerat, S., Yildiz, A. R. & Mirjalili, S. A comparative study of recent multi-objective metaheuristics for solving constrained truss optimisation problems. Arch. Comput. Methods Eng. 28, 4031– 4047 (2021).

38 Das, R. & Baker, D. Macromolecular modeling with rosetta. Annu. Rev. Biochem. 77, 363–382 (2008).

39 Guest, J. D. et al. An expanded benchmark for antibody-antigen docking and affinity prediction reveals insights into antibody recognition determinants. Structure 29, 606–621 (2021).

40 Berman, H. M. et al. The protein data bank. Nucleic Acids Res. 28, 235–242 (2000).

41 Dunbar, J. et al. SAbDab: the structural antibody database. Nucleic Acids Res. 42, D1140–D1146 (2014).

42 Ruffolo, J. A., Chu, L.-S., Mahajan, S. P. & Gray, J. J. Fast, accurate antibody structure prediction from deep learning on massive set of natural antibodies. Nat. Commun. 14, 2389 (2023).

43 Basu, S. & Wallner, B. DockQ: a quality measure for protein-protein docking models. PloS One 11, e0161879 (2016).

44 Zhang, Y. & Skolnick, J. Scoring function for automated assessment of protein structure template quality. *Proteins Struct., Funct.*, Bioinf. 57, 702–710 (2004).

45 Fadini, A. et al. Highlights of model quality assessment in CASP16. *Proteins Struct., Funct.*, Bioinf. (2025).

46 Guo, Z., Liu, J., Skolnick, J. & Cheng, J. Prediction of inter-chain distance maps of protein complexes with 2D attention-based deep neural networks. Nat. Commun. 13, 6963 (2022).

47 Xu, Y., Xu, D. & Gabow, H. N. Protein domain decomposition using a graph-theoretic approach. Bioinformatics 16, 1091–1104 (2000).

48 Steinegger, M. & Söding, J. MMseqs2 enables sensitive protein sequence searching for the analysis of massive data sets. Nat. Biotechnol. 35, 1026–1028 (2017).

49 Lefranc, M.-P. et al. IMGT, the international ImMunoGeneTics database. Nucleic Acids Res. 27, 209–212 (1999).

50 Hou, M. et al. High-accuracy protein complex structure modeling based on sequence-derived structure complementarity. Preprint at bioRxiv, 10.1101/2025.03.26.645390 (2025).

51 Mirdita, M. et al. Uniclust databases of clustered and deeply annotated protein sequences and alignments. Nucleic Acids Res. 45, D170–D176 (2017).

52 Remmert, M., Biegert, A., Hauser, A. & Söding, J. HHblits: lightning-fast iterative protein sequence searching by HMM-HMM alignment. Nat. Methods 9, 173–175 (2012).

53 Lin, T.-Y., Goyal, P., Girshick, R., He, K. & Dollár, P. Focal loss for dense object detection. Proceedings of the IEEE international conference on computer vision, 2980–2988 (2017).

54 Silverman, B. W. Density estimation for statistics and data analysis. Routledge, (2018).

## Supplementary References

1 Abramson, J. et al. Accurate structure prediction of biomolecular interactions with AlphaFold 3. Nature 630, 493–500 (2024).

2 Evans, R. et al. Protein complex prediction with AlphaFold-Multimer. Preprint at biorxiv, 10.1101/2021.10.04.463034 (2021).

3 Yan, Y., Tao, H., He, J. & Huang, S.-Y. The HDOCK server for integrated protein–protein docking. Nat. Protoc. 15, 1829–1852 (2020).

4 Guest, J. D. et al. An expanded benchmark for antibody-antigen docking and affinity prediction reveals insights into antibody recognition determinants. Structure 29, 606–621.e605 (2021).

5 Olechnovič, K. & Venclovas, Č. VoroIF-GNN: Voronoi tessellation-derived protein–protein interface assessment using a graph neural network. *Proteins Struct., Funct.*, Bioinf. 91, 1879–1888 (2023).

6 Storn, R. & Price, K. Differential evolution–a simple and efficient heuristic for global optimization over continuous spaces. J. Glob. Optim. 11, 341–359 (1997).

7 Meng, Z. et al. Application of state-of-the-art multiobjective metaheuristic algorithms in reliability-based design optimization: a comparative study. Struct. Multidiscip. Optim. 66, 191 (2023).

8 Panagant, N., Pholdee, N., Bureerat, S., Yildiz, A. R. & Mirjalili, S. A comparative study of recent multi-objective metaheuristics for solving constrained truss optimisation problems. Arch. Comput. Methods Eng. 28 (2021).

9 Das, R. & Baker, D. Macromolecular modeling with rosetta. Annu. Rev. Biochem. 77, 363–382 (2008).

10 Dunbar, J. & Deane, C. M. ANARCI: antigen receptor numbering and receptor classification. Bioinformatics 32, 298–300 (2016).

11 Lefranc, M.-P. et al. IMGT, the international ImMunoGeneTics database. Nucleic Acids Res. 27, 209–212 (1999).

12 Basu, S. & Wallner, B. DockQ: a quality measure for protein-protein docking models. PloS One 11, e0161879 (2016).

13 Zhang, Y. & Skolnick, J. Scoring function for automated assessment of protein structure template quality. *Proteins Struct., Funct.*, Bioinf. 57, 702–710 (2004).

14 Meiler, J., Müller, M., Zeidler, A. & Schmäschke, F. Generation and evaluation of dimension-reduced amino acid parameter representations by artificial neural networks. Molecular Modeling Annual 7, 360–369 (2001).

15 Rao, R. et al. MSA Transformer. Proceedings of the 38th International Conference on Machine Learning. 139, 8844–8856 (2021).

16 Ruffolo, J. A., Gray, J. J. & Sulam, J. Deciphering antibody affinity maturation with language models and weakly supervised learning. Preprint at arXiv, 10.48550/arXiv.2112.07782 (2021).

17 Altschul, S. F. et al. Gapped BLAST and PSI-BLAST: a new generation of protein database search programs. Nucleic Acids Res. 25, 3389–3402 (1997).

18 Ballester, P. J. & Richards, W. G. Ultrafast shape recognition to search compound databases for similar molecular shapes. J. Comput. Chem. 28, 1711–1723 (2007).

19 Krapp, L. F., Abriata, L. A., Cortés Rodriguez, F. & Dal Peraro, M. PeSTo: parameter-free geometric deep learning for accurate prediction of protein binding interfaces. Nat. Commun. 14, 2175 (2023).

20 Remmert, M., Biegert, A., Hauser, A. & Söding, J. HHblits: lightning-fast iterative protein sequence searching by HMM-HMM alignment. Nat. Methods 9, 173–175 (2012).

21 Zhang, Y., Zhou, D., Chen, S., Gao, S. & Ma, Y. Single-image crowd counting via multi-column convolutional neural network. Proceedings of the IEEE conference on computer vision and pattern recognition, 589–597 (2016).

22 Lin, P., Tao, H., Li, H. & Huang, S.-Y. Protein–protein contact prediction by geometric triangle-aware protein language models. *Nat*. Mach. Intell. 5, 1275–1284 (2023).

